# Activation of Autoreactive Lymphocytes in the Lung by STING Gain-of-function Mutation Radioresistant Cells

**DOI:** 10.1101/2023.07.28.551002

**Authors:** Kevin MingJie Gao, Kerstin Nündel, Kristy Chiang, Xihui Yin, Paul J. Utz, Kate Fitzgerald, Ann Marshak-Rothstein

## Abstract

Gain-of-function mutations in the dsDNA sensing adaptor STING lead to a severe autoinflammatory syndrome known as STING-associated vasculopathy with onset in Infancy (SAVI). SAVI patients develop interstitial lung disease (ILD) and commonly produce anti-nuclear antibodies (ANAs), indicative of concomitant autoimmunity. Mice heterozygous for the most common SAVI mutation, V154M (VM), also develop ILD, triggered by nonhematopoietic VM cells, but exhibit severe peripheral lymphopenia, low serum Ig titers and fail to produce autoantibodies. In contrast, we now show that lethally irradiated VM mice reconstituted with WT stem cells (WT→VM chimeras) develop ANAs and lung-reactive autoantibodies associated with accumulation of activated lymphocytes and formation of germinal centers in lung tissues. Moreover, when splenocytes from WT→VM chimeras were adoptively transferred into unmanipulated Rag1^-/-^ mice, donor T cells accumulated in the lung. Overall, these findings demonstrate that expression of the VM mutation in non-hematopoietic cells can promote the activation of immunocompetent autoreactive lymphocytes.

**Summary:** Chimeric mice expressing STING only in non-hematopoietic cells develop systemic and lung directed autoimmunity which recapitulates what is seen in pediatric patients with SAVI disease.

## Introduction

Autoinflammatory and autoimmune diseases exist on a continuum, reflecting the connection between innate and adaptive immunity. Both can involve dysregulation of immune responses that leads to a breach in tolerance and the ensuing activation of autoreactive lymphocytes. While some diseases may originate entirely from the dysregulation of adaptive immune responses^1^, many diseases commonly recognized as autoimmune can be triggered by aberrant activation of innate immune processes. For example, in the autoimmune disease Systemic Lupus Erythematosus (SLE), inadequate clearance of cell debris in combination with activation of endosomal Toll-like receptors leads to the activation of B cells reactive to nucleic acids (**NA**) or NA-binding proteins^2^. Similar autoantibody reactivities develop in patients with a range of autoinflammatory diseases that result from defects in nucleic acid sensing, degradation, or metabolism^3–5^. One such autoinflammatory disease is SAVI, a syndrome caused by constitutive ligand-independent activation of STING, the adaptor protein in the cGAS-STING cytosolic dsDNA sensing pathway. SAVI patients commonly develop interstitial lung disease (ILD), and in a cohort of 21 SAVI patients, 20 were found to be positive for autoantibodies^5^.

Mouse models of SAVI that express the two most common SAVI causing STING mutations, STING N153S (NS) and STING V154M (VM), recapitulate many of the clinical manifestations found in patient populations, including ILD^6, 7^. VM mice develop more severe ILD and, despite the accumulation of activated extravascular B cells in the lung, they also develop peripheral B cell lymphopenia much earlier in life than NS mice^8, 9^, and exhibit a more severe defect in VM B cell development and antibody production. Expression of the VM mutation in T cells is known to compromise T cell survival by directly triggering an unfolded protein response (UPR) that causes apoptotic cell death ^10, 11^. Analogously, STING activation in B cells has also been found to impair B cell survival^9, 12^. In contrast to mouse B cells, human B cells do not express STING^13^ and human SAVI patients are not B cell lymphopenic.

We previously reported that lethally irradiated VM mice reconstituted with WT stem cells (WT→VM radiation chimeras) are not lymphopenic and develop more extensive lung inflammation than the parental VM mice^14^. These data indicate that non-hematopoietic cells expressing the VM mutation can recruit and activate WT lymphocytes and thereby play a significant role in triggering ILD. Since lymphocyte-intrinsic STING GOF is the primary driver of T and B cell lymphopenia^6^, we decided to use non-lymphopenic WT→VM chimeras^15^ to explore the impact of the VM host environment on potentially autoreactive WT lymphocytes. We reasoned that these chimeras would better model the autoimmune phenotype of SAVI patients. We further incorporated BCR and TCR transgenic mice into our model to assess the importance of self-reactivity in lymphocyte recruitment and activation in the lung. We now show that WT autoreactive B cells in a VM host become activated and produce autoantibodies reactive with common autoantigens as well as with lung tissue. We further found that the VM host promoted the activation of potentially self-reactive T cells and the development of long-lived lung-homing T cell subset. Altogether, our findings demonstrate that autoinflammation resulting from STING GOF in non-hematopoietic cells triggers both B cell and T cell autoimmunity.

## Results

### B cell intrinsic STING GOF impairs proliferation and antibody production

VM SAVI mice are lymphopenic and have been reported to have remarkably low serum antibody titers^9^. Consistent with these observations, we found that VM serum IgM, IgG1, and IgG2c titers were reduced by approximately 10-fold, 100-fold, and 1000-fold, respectively, relative to littermate controls (**Figure 1A**). Low antibody titers could reflect B cell intrinsic defects in survival, proliferation, and/or differentiation or they could reflect the impact of B cell extrinsic factors. Because lethally irradiated WT hosts reconstituted with VM BM poorly engraft donor B cells^8^, it was not possible to directly assess the effects of BM intrinsic VM expression on B cell phenotypes. Instead, we asked whether hypogammaglobulinemia was due to the VM environment by lethally irradiating VM or WT mice and reconstituting them with WT stem cells (WT➔VM and WT➔WT radiation chimeras). 8-10 weeks later, serum Ig titers collected from WT➔VM chimeras were compared to serum titers from WT➔WT chimera controls. IgM and IgG1 titers were comparable, while IgG2c titers were only reduced by 10-fold (**Figure 1A**). These data indicate that low Ab titers of VM mice are not attributable to the VM environment and most likely due to a B cell intrinsic defect, although the host environment may further contribute to the loss of IgG2c-producing cells. We speculated that low antibody titers could reflect a STING-dependent ER stress response of VM B cells following BCR engagement^10, 12^ that impairs B cell survival and proliferation.

**Figure 1.**
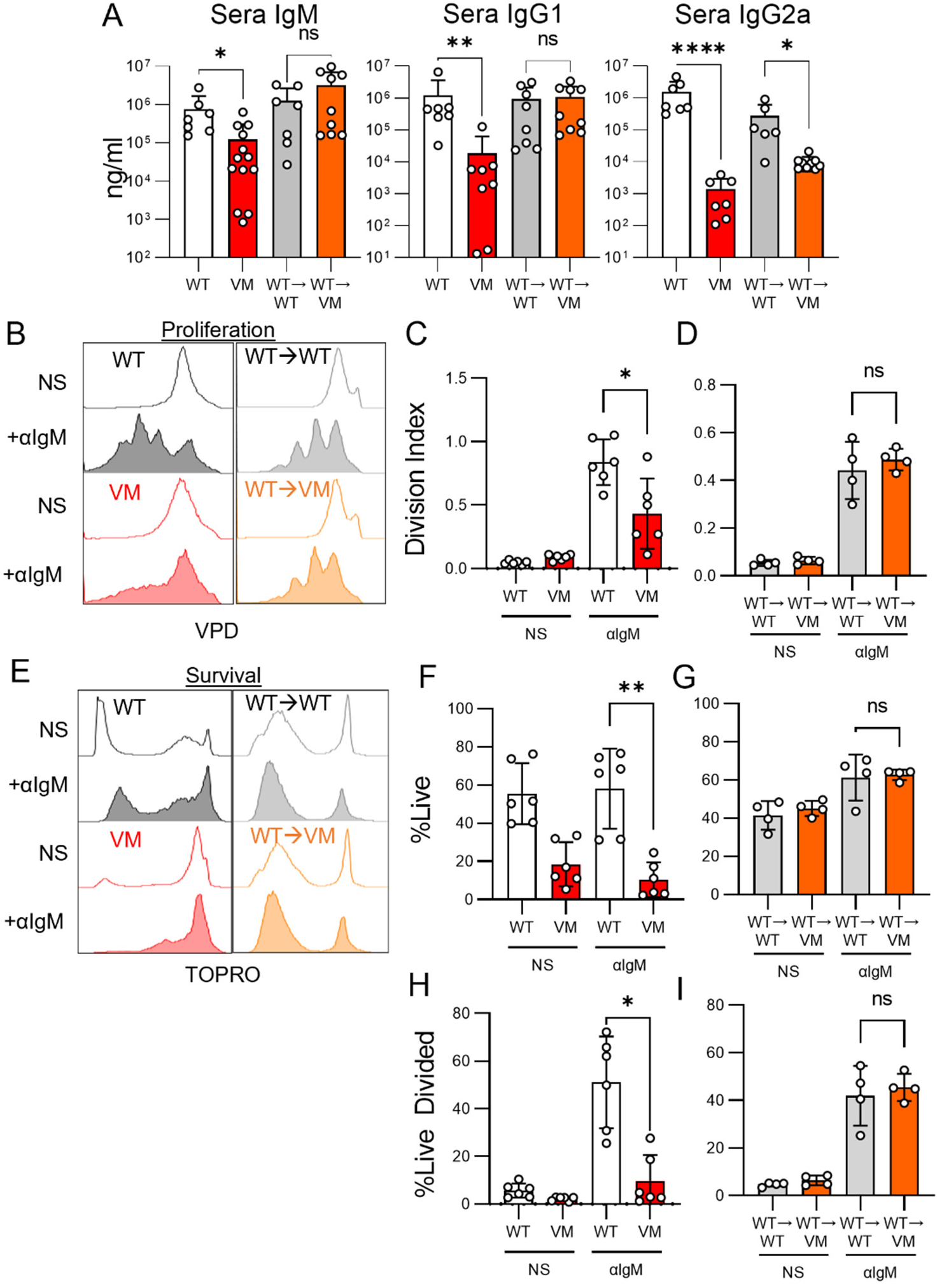
Reduced antibody production, B cell proliferation and B cell survival reflect B cell intrinsic STING gain-of-function. (**A**) Serum titers of IgM, IgG1 and IgG2a in age- and sex-matched WT (n=7) and VM (n=7-14) mice at 3-4 mons of age, or from WT→WT (n=7) and WT→VM (n=9) chimeric mice at 8 weeks post reconstitution. (**B-I**) Splenic B220+ B cells from biologic replicates of WT (n=6) vs VM (n=6) B cells, and WT→WT (n=4) and WT→VM (n=4) B cells were isolated by magnetic bead purification and labelled with violet proliferative dye (VPD) prior to a 72-hour stimulation with BLyS (NS) alone or anti-IgM F(ab’)2 antibody and BLyS (+αIgM). Cells were stained with TOPRO immediately prior to analysis to assess cell death. (**B**) Representative histograms of VPD intensity as a metric of cell division (**C-D**) Division index (average divisions per cell from original population) (**E**) shows representative histograms of TOPRO intensity as a metric of cell death in B cells. (**F-G**) Percentage of live cells as defined as TOPRO negative (**H-I**) Percentage of live divided cells as defined as TOPRO negative VPD lo. Nonparametric Mann-Whitney U-tests were used for pair-wise comparisons, and a non-parametric Kruskal-Wallis test was used for one-way ANOVA to determine statistical significance (ns p>0.05, *p<0.05,**p<0.01, ***p<0.001, ****p<0.0001).

To test this hypothesis, we compared the in vitro survival and proliferative capacity of parental WT and VM B cells to B cells isolated from the chimeric mice. Splenic B cells were stained with the fluorophore VPD, and then stimulated with anti-IgM F(ab’)_2_. 3 days post-stimulation, proliferation was assessed by VPD dilution and viability was evaluated with TOP-RO-3 (dead cell) stain. As expected, WT B cells proliferated well in response to anti-IgM, whereas the VM B cells proliferated poorly (**Figure 1B,C**) and had reduced survival, even in the presence of the B cell survival factor BLyS (**Figure 1E,F**). Based on these observations, we speculate that VM B cells show an activation induced demise precluding robust proliferation. Consistently the fraction of live divided B cells is significantly attenuated in VM compared to WT B cells following activation (Figure 1H). This data is also in agreement with previous studies of STING agonists in B cells^12^, and analogous to the increased cell death observed in SAVI T cells following TCR engagement^10^. By contrast, we found that B cells from the WT➔VM chimeras responded comparably to control WT➔WT B cells (**Figure 1B,D,E,G,I**). Thus, B cells isolated from WT➔VM chimeras are unimpaired as far as their capacity to proliferate and differentiate, despite the ongoing systemic inflammation and the robust accumulation of B cells in the lungs of these mice^14^. Therefore, we surmised that WT➔VM chimeras provide a reasonable model for exploring the recruitment of potentially autoreactive B cells to the lung in the context of SAVI ILD.

### WT→VM chimeric mice develop germinal centers in the lung

Germinal centers (GC) are a common site of B cell expansion, differentiation, and class switch recombination^16^. Unimmunized WT mice spontaneously develop small numbers of GC in the spleen and have low but detectable numbers of Fas+ GL7+ GC splenic B cells^17^. VM mice have fewer B cells than WT mice^6^ and the percentage of GC B cells within the total B cell compartment is also reduced (**Figure 2A**). This is in line with the impaired in vitro responses of VM B cells described above. In contrast, the percentage of GC B cells in both the WT→WT and WT→VM chimeras was significantly higher than that of VM mice. Several of the WT→VM chimeric mice even had notably higher percentages of GC B cells (p=0.2283).

**Figure 2.**
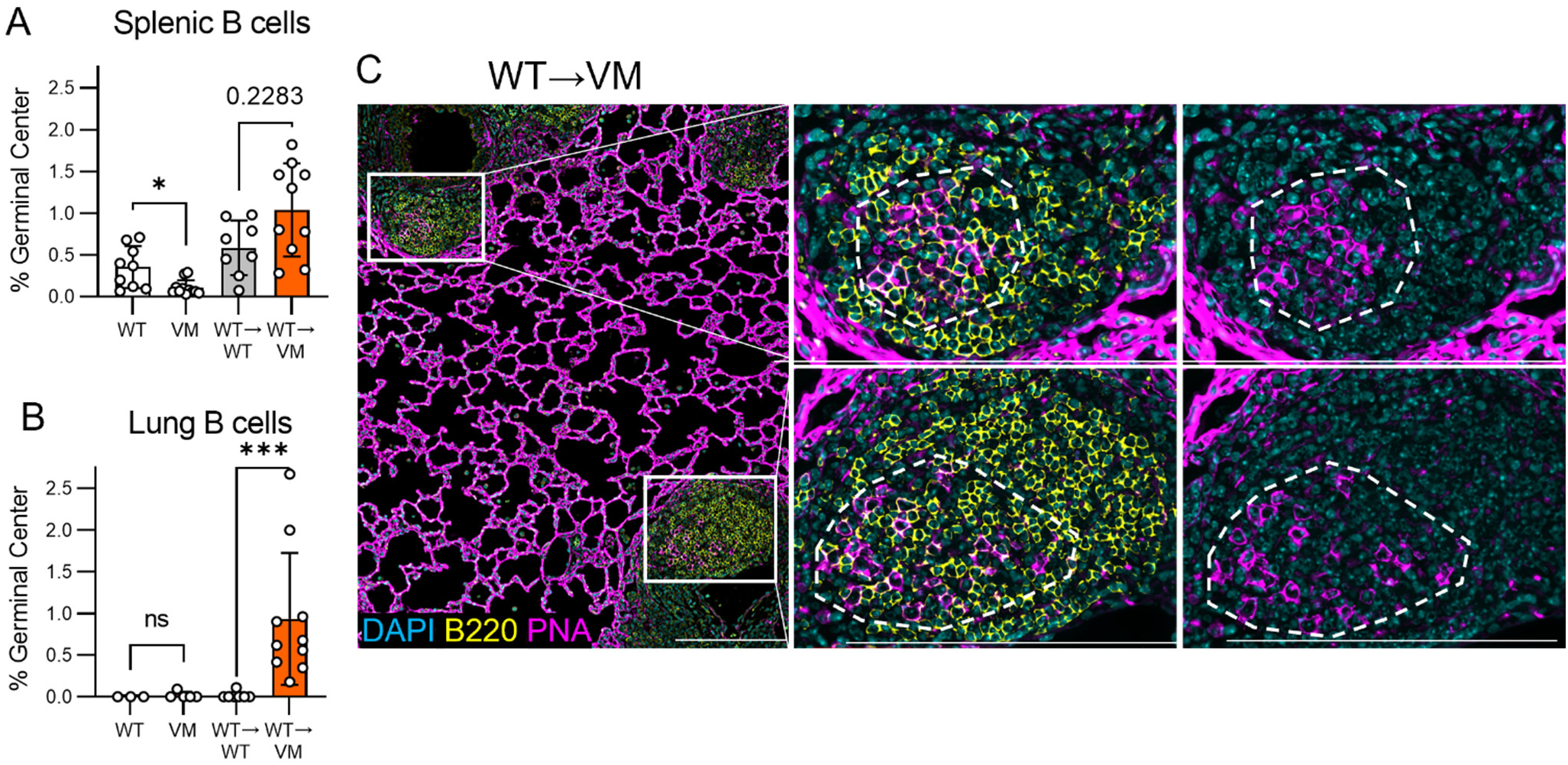
WT→VM mice form germinal centers in the lung. Data obtained from 3-4 month-old age- and sex-matched WT (n=9) and VM (n=11) mice. Chimeric WT→WT (n=8) and WT→VM (n=10) mice were evaluated 8 to 9 weeks post-reconstitution. (**A**) Percentage of IgD-Fas+ GL7+ germinal center (GC) B cells within CD45+ CD19+ B cell compartment of WT and VM spleen, or within the donor-derived B cells in WT→WT and WT→VM chimeric spleens. (**B**) Percentage of GC B cells within the CD45+ CD19+ lung extravascular (EV) B cell compartment of in WT and VM mice, and within the total donor-derived lung EV B cells in WT→WT and WT→VM chimeras. (**C**) Immunofluorescence imaging of WT→VM chimeric mouse lungs stained for B220 (yellow) and peanut agglutinin (PNA, magenta) to identify B220+ PNA+ GC B cells. Two B cell follicles containing GCs are highlighted with white squares. 3x magnification of the highlighted sections are shown in the 2 panels to the right, and a dotted circular outline is used to identify the germinal center. In the middle panel, DAPI, B220, and PNA are shown together, and the right panel shows DAPI and PNA alone. A non-parametric Kruskal-Wallis test was used for one-way ANOVA to determine statistical significance (ns p>0.05, *p<0.05, ***p<0.001). Nonparametric Mann-Whitney U tests were used for pairwise comparisons to determine statistical significance (**p<0.01, ***p<0.001). Scale bars in C represent 200µm.

Well-organized bronchus associated lymphoid tissue (BALT) can be found in the inflamed lungs of patients with autoimmune diseases such as rheumatoid arthritis and Sjogren’s Syndrome and can include follicles with defined GCs^18^. GCs within the lungs of WT→VM chimeric mice were visualized by immunofluorescent staining for peanut agglutinin (PNA), a lectin that binds to GC B cells. Although PNA also binds lectin receptors on alveolar epithelia, PNA+ GC cells could be identified within CD45+ B220+ B cell follicles (**Figure 2C**). Although both VM and WT→VM chimeric mice exhibited extensive BALT^14^, only BALT from WT→VM chimeric mice contained a significant number of GC B cells (**Figure 2B**). Thus, immunocompetent (WT) B cells can be induced to form GC in the lungs of chimeric mice where radioresistant cells express the VM mutation.

### Activation of EV B cells requires a diverse repertoire

BALT and GCs are normally associated with antigen-driven expansion of pathogen or autoantigen-specific B cells. Based on the frequency of GCs in the WT→VM chimeric SAVI mice, we hypothesized that a STING GOF lung environment recruits and then activates autoreactive lymphocytes. Alternatively, lymphocytes could theoretically be non-specifically recruited and activated in a repertoire independent fashion by the inflammatory milieu present within the SAVI lung. We reasoned that mixed radiation chimeras would facilitate our ability to track the relative accumulation of repertoire restricted (BCR transgenic (Tg)) and polyclonal (non-Tg) B cells in the lungs of VM host mice.

A mixture of CD45 allelically distinct BM stem cells from Rag1^-/-^ BCR Tg MD4 (hen egg lysozyme-specific) mice and non-Tg B6 mice at a 4:1 ratio was used to reconstitute either lethally irradiated WT mice (MD4+WT→WT chimeras) or VM mice (MD4+WT→VM chimeras), and mice were evaluated 8 weeks later (**Figure 3A**). The capacity of the WT- and MD4-derived stem cells to engraft the overall hematopoietic compartment was determined by examining the relative proportion of the donor populations in the BM of the chimeric mice. As expected, the composition of the hematopoietic compartment in the BM reflected the initial injection ratio of 80% MD4/20% WT in both the WT and VM recipients (**Figure 3B**). In the spleen and extravascular (EV) lung, the percentage of MD4-derived B cells was reduced to ∼ 20%, as might be anticipated for cells with a restricted repertoire (**Figure 3B**). However, the total number of both WT and MD4-derived donor B cells in the EV lung was, as expected, much higher in VM recipients (**Figure 3C**). Together, these data show that the non-Tg WT B cells generally outcompeted Tg MD4 B cells as far as accumulation in peripheral lymphoid compartments. However, MD4 cells were not preferentially excluded from the lungs of VM recipients. Therefore, recruitment of B cells to STING GOF lung appears to be repertoire independent. In contrast, only WT donor cells in the lungs of the VM host became Fas^+^ GL7^+^ GC B cells; whereas, MD4 donor cells in VM host lungs did not become GC B cells (**Figure 3D**). Thus, the formation of lung GCs requires a diverse repertoire that includes potentially autoreactive B cells.

**Figure 3.**
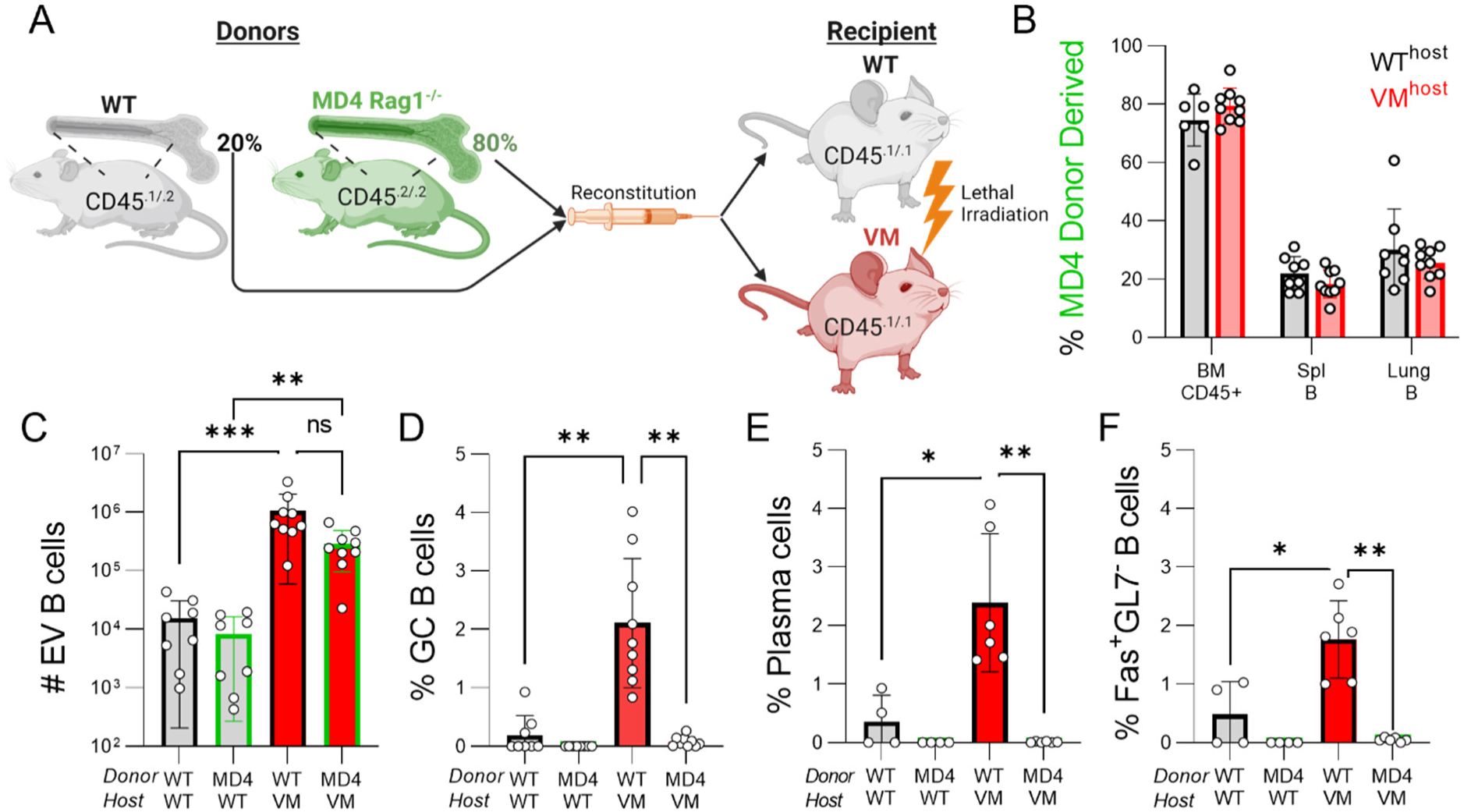
Formation of GC B cells in VM lung requires an unrestricted repertoire. (**A**) 6-week-old age- and sex-matched CD45.1/1 WT and VM littermates were lethally irradiated and reconstituted with a mixture of bone marrow stem cells comprised of 20% CD45.1/2 WT and 80% CD45.2/2 MD4 Rag1^-/-^ cells. WT+MD4➔WT (n=4-8) and WT+MD4➔VM (n=6-9) chimeric mice were then evaluated 8 to 9 weeks later. (**B**) percentage of MD4 donor-derived cells within all CD45+ cells in bone marrow (BM), within the splenic CD19+ B compartment, or within the lung extravascular (EV) B cell compartment of either the WT hosts (grey bar) or VM hosts (red bar), after excluding any CD45.1 single positive radio-resistant host cells. (**C**) total number of lung EV B cells derived from either the WT or MD4 donor in either the WT or VM hosts. (**D**) percentage of IgD^neg^ Fas^+^ GL7^+^ germinal center B cells derived from either the WT or MD4 donor within the EV B cell compartment of either WT or VM host. (**E**) percentage of B220^neg^ CD138^+^ PC cells derived from either the WT or MD4 donor within the EV B cell compartment of either WT or VM host from WT donor and MD4 donor-derived lung EV B cells in WT and VM hosts. (**F**) percentage of IgM^+^ IgD^neg^ Fas^+^ GL7^neg^ B cells derived from WT or MD4 donor within the EV B compartment of WT or VM hosts. (**B-E**) Each dot represents an individual mouse. Nonparametric Mann-Whitney U-tests were used for pair-wise comparisons, and a non-parametric Kruskal-Wallis test was used for one-way ANOVA to determine statistical significance (ns p>0.05, *p<0.05, **p<0.01, ***p<0.001).

Additional flow cytometric analysis also revealed a significant enrichment of WT-derived plasma cells (PCs) among the lung B cells of the VM but not WT recipients, whereas MD4+ PCs were not detected in either host (**Figure 3E**). We also found a small population of non-Tg Fas+ CD86+ GL7-IgM+ IgD-cells enriched in the VM recipients but not the WT recipients; this phenotype is associated with Age-associated B cells (ABCs)^19, 20^ (**Figure 3F**). Thus, in addition to the repertoire dependent generation of GC B cells in the VM host, the VM host environment also favors other subsets potentially associated with autoantibody production.

Beyond the specific subsets identified by conventional flow cytometric analysis, multiparameter flow cytometry can be evaluated through dimensional reduction (**Figure S1A**), unsupervised clustering and manual cluster annotation of multiparameter flow cytometry data as a complementary analytic approach to visualize broader patterns B cell differentiation (**Figure S1B, Table 1**). The UMAP of our lung B cell data was organized such that UMAP1^hi^ (x-axis) was associated with IgD and UMAP2^hi^ (y-axis) was associated with IgM and CD69 (**Figure S1C**). Our analysis affirmed that the B cell subsets identified in Fig. 3, GC B cells (green oval), PCs (black oval), and ABC-like B cells (red circle), were uniquely elevated in our WT→VM B cells (Figure S1D). UMAP further revealed a dramatic difference in the distribution of non-Tg and MD4 cells in the WT hosts, where MD4 B cells were more skewed towards an IgM^hi^, IgD^lo^, CD69^hi^ phenotype, indicative of a more activated population. By contrast, in the VM recipient lungs, apart from the PC, GC and ABC clusters, the overall distribution of the majority of the EV non-Tg and MD4 cells is much more similar and can be considered IgM^lo^ IgD^hi^, consistent with a less activated or potentially anergic phenotype^21^. The MD4 B cells in the VM recipients express somewhat higher levels of IgM than the non-Tg B cells, consistent with what is found in MD4 Tg mice^22^. This may reflect repertoire independent features of the VM lung environment which attract naïve B cells and/or overall alterations in the circulating B cell pool within a VM host. Collectively, we find that the VM lung effectively recruits both non-Tg and WT naive B cells that require a diverse repertoire to become further activated.

**Table 1.**
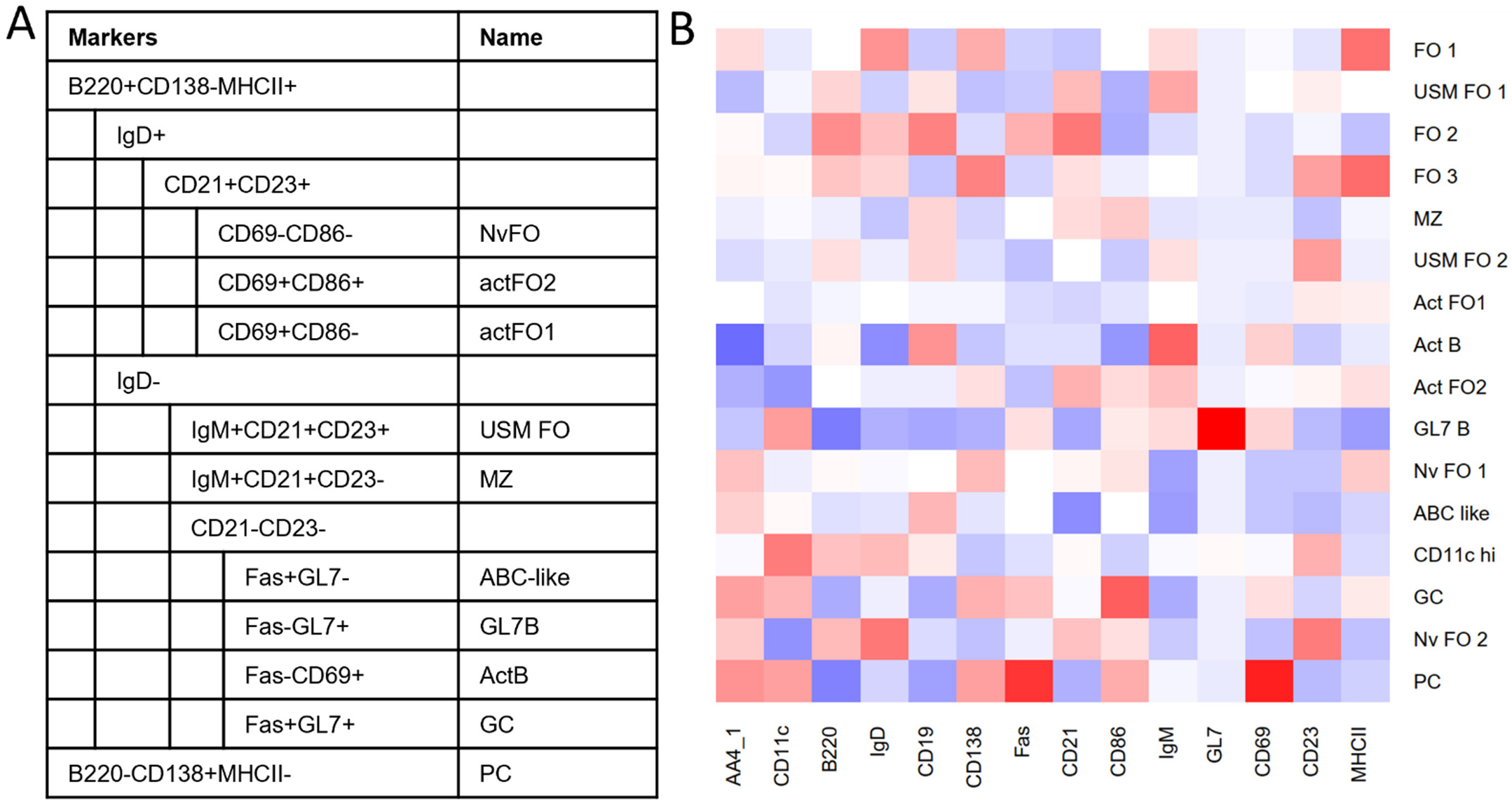
Annotation of B cell clusters by marker expression. (A) Manual annotation of 10 clusters identified by Phenograph unsupervised clustering of Live CD45+CD45IV-CD19+ lung extravascular (EV) B cell flow cytometry events from WT+MD4→WT (n=4) and WT+MD4→VM (n=6) mice concatenated after excluding CD45.1 single positive radioresistant events.(B) shows a column normalized heatmap of median marker expression for clusters described in (A) and Figure S1B. Blue represents lower expression, white represents intermediate expression, red indicates higher expression.

**Figure S1.**
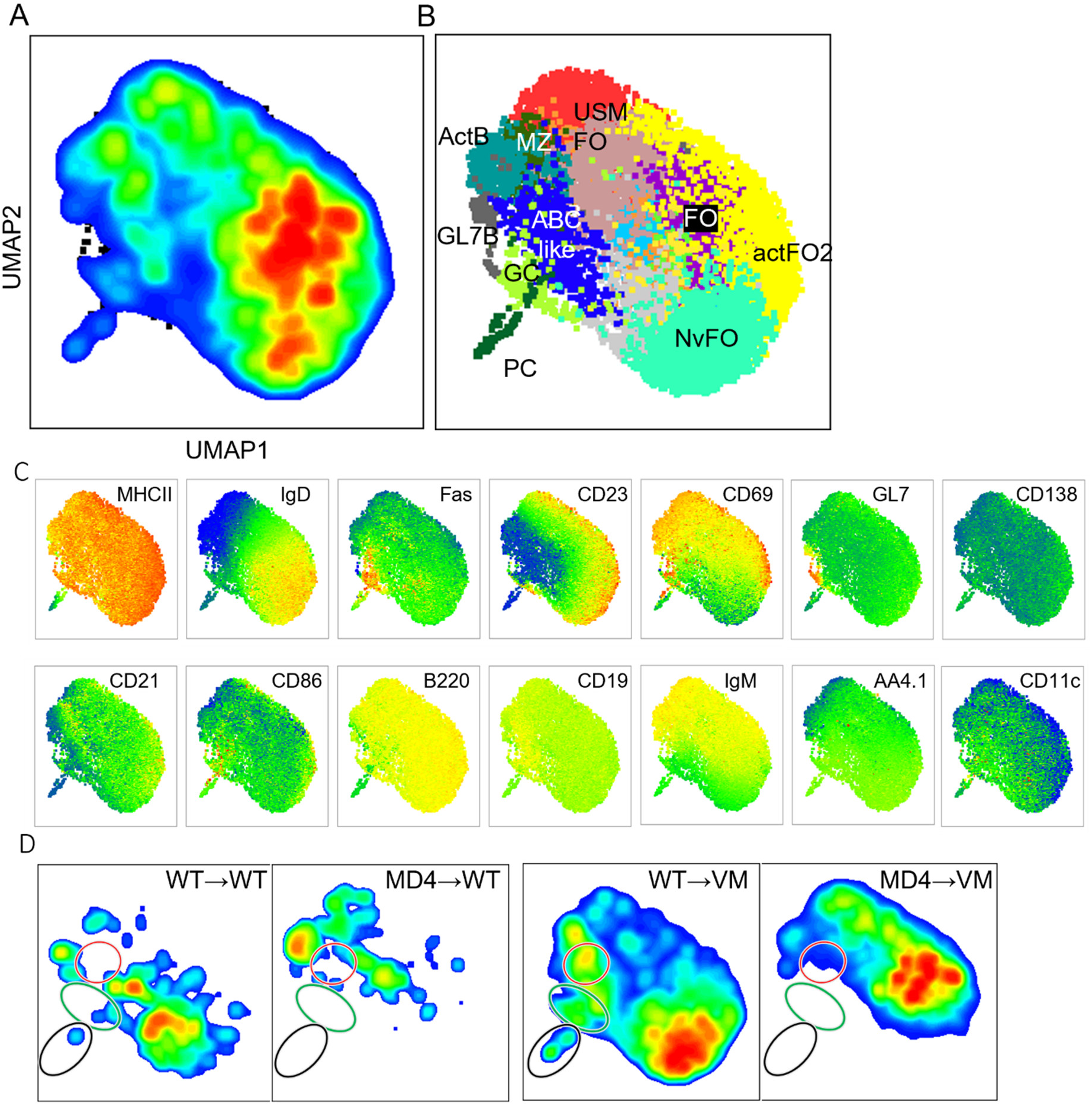
Unsupervised clustering of B cell markers identifies repertoire and VM host dependent phenotypes. 1000 Live CD45+CD45IV-CD19+ lung extravascular (EV) B cell flow cytometry events per mouse from WT+MD4→WT (n=4) and WT+MD4→VM (n=6) mice were concatenated after excluding CD45.1 single positive radioresistant events. (**A**) A UMAP was generated using MHCII, IgD, Fas, CD23, CD69, GL7, CD138, CD21, CD86, B220, CD19, IgM, AA4.1, and CD11c. (**B**) Unsupervised clustering using Phenograph identified 10 clusters in this dataset, which were manually annotated based on their expression profile (Table 1) and projected onto the UMAP alongside their annotated name. (**C**) Heatmap projections for each parameter used to generate the UMAP and perform unsupervised clustering. (**D**) UMAPs depicting events from WT donor-derived cells in either WT (WT→WT) or VM (WT→VM) hosts, as well as events from MD4 donor-derived cells in either WT (MD4→WT) or VM (MD4→WT) hosts. Regions of the UMAP enriched in WT→VM events are highlighted with rings corresponding to the ABC-like cluster (red), the germinal center (GC) cluster (green), and the plasma cell (PC) cluster (black).

### Non-Transgenic T cells are preferentially recruited to and activated in the VM lung

TCRαβ T cell infiltration of the lung is a predominate feature of SAVI disease, and as with B cells, WT T cells are recruited and activated in the lungs of VM hosts^14^. To determine whether a restricted TCR repertoire limits recruitment and/or activation of T cells to the SAVI lung, we utilized OTI and OTII transgenic mice which express TCRs specific for ovalbumin peptides bound to MHCI and MHCII, respectively. OTI T cells become CD8 T cells, whereas OTII T cells become CD4 T cells. Radiation chimeras were again used to assess the dependence of repertoire on T cell recruitment to the lungs in both CD4 and CD8 T cell subsets. To reconstitute both the CD4 and CD8 host compartments with repertoire restricted donors, lethally irradiated WT or VM hosts were given a mixture of non-Tg WT (20%) and allelically distinct Rag2^-/-^ OTI (40%) and Rag2^-/-^ OTII (40%) stem cells (1:2:2 ratio), indicated as OT+WT➔WT and OT+WT➔VM, respectively (**Figure 4A**). We then examined the BM, spleen, and lungs of chimeric mice 8 weeks later.

**Figure 4.**
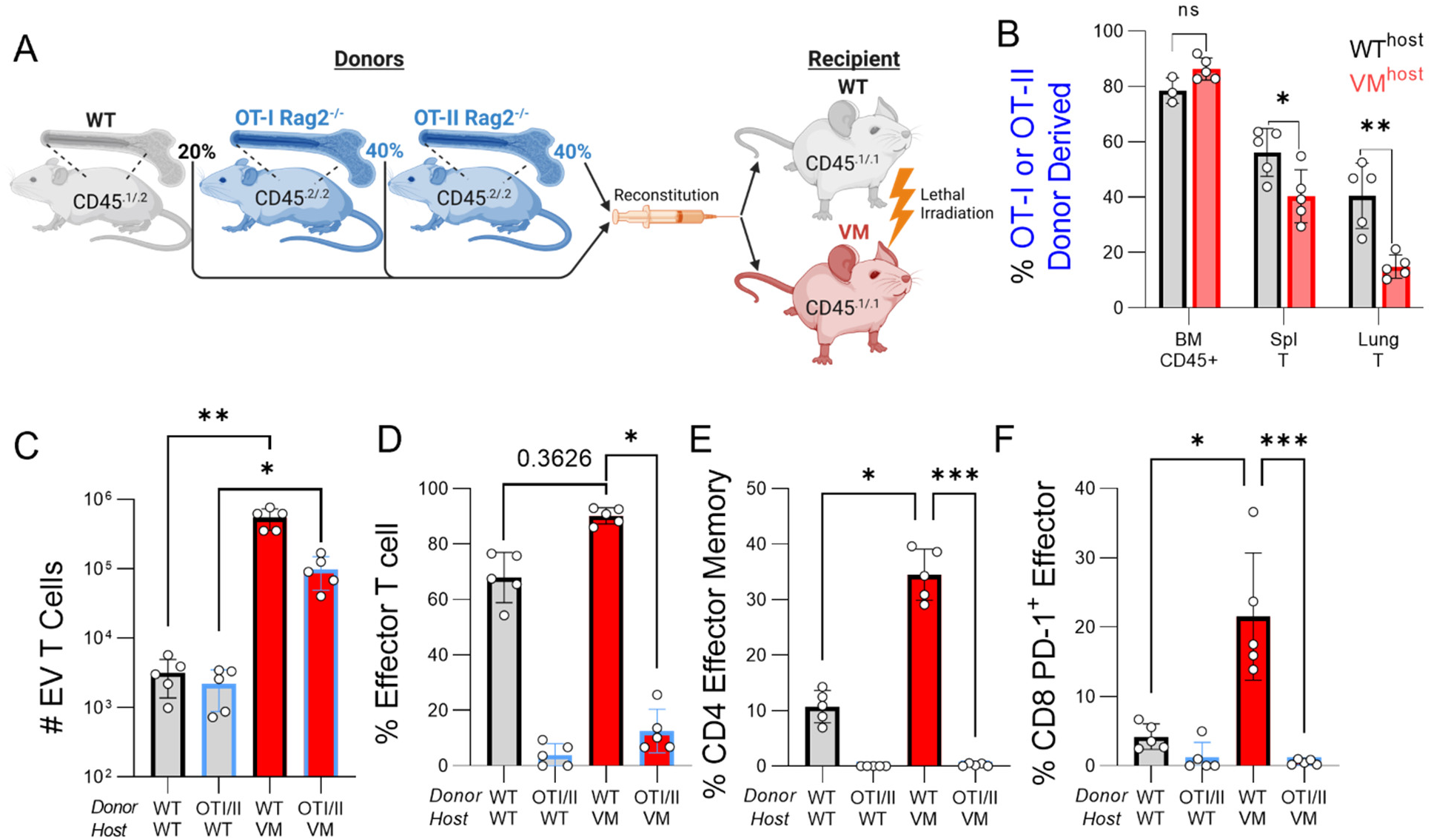
T lymphocyte recruitment and activation in VM ILD requires a diverse repertoire. (**A**) 6-week-old sex-matched CD45.1/1 WT and VM littermates were lethally irradiated and reconstituted with a mixture of BM stem cells comprised of 20% WT CD45.1/2, 40% OT-I Rag2^-/-^ CD45.2/2, and 40% OT-II Rag2^-/-^ CD45.2/2 donor bone marrow. WT+OTI/II→WT (n=5) and WT+OTI/II→VM (n=5) chimeric mice were evaluated 8 to 9 weeks later. (**B**) % OTI/II donor-derived cells within the donor-derived total BM compartment, CD3+ splenic T cell compartment, and CD3+ lung extravascular (EV) T cell compartment. (**C**) Total number of lung EV T cells derived from either the WT donor or the OTI/II donors in WT and VM hosts. (**D**) % CD44+CD62L-effector T cells within the WT and MD4 donor-derived lung EV T cell compartment in WT and VM hosts. (**E**) % CD127+ CD69-CD4 Eff Mem cells within the WT and OTI/II donor-derived lung EV ab CD4 T cell compartment in WT and VM hosts. (**F**) % PD-1+ CD69+ CD8 Eff PD1 cells within the WT and OTI/II donor-derived lung EV ab CD8 T cell compartment in WT and VM hosts. A non-parametric Kruskal-Wallis test was used for one-way ANOVA to determine statistical significance (ns p>0.05, *p<0.05,**p<0.01, ***p<0.001).

As in the BCR Tg experiments, the percentage of TCR Tg (OTI + OTII) donor-derived CD45+ BM cells reflected the donor cell inoculum (∼80%) in both WT and VM hosts (**Figure 4B**). However, in the spleen, the frequency of Tg T cells in the VM hosts (∼40%) was reduced compared to the WT hosts (∼60%). The differences in the lung EV B cell compartment were even more dramatic as 40% of T cells in the lungs of WT hosts were derived from OT Tg donors, while only 10% of T cells were derived from OT Tg donors in the lungs of VM hosts (Figure 4B). However, consistent with the increased number of CD45+ EV T cells in VM mice, both OT and WT donor-derived T cells accumulated at significantly higher numbers in the VM host (**Figure 4C**). Overall, OT donor T cells were significantly outcompeted by repertoire unrestricted WT donor T cells in spleen and lungs of VM hosts. These data indicate that a diverse repertoire improves T cell recruitment to VM lung.

Moreover, CD44+CD62L-effector subsets were almost entirely derived from the WT donor (**Figure 4D**), indicating that a diverse TCR repertoire favored T cell activation within the lung. There was a somewhat higher percentage of effector cells within the WT T cell compartment of the VM host compared to WT hosts (p=0.3626). To further identify differences in effector T cell phenotypes arising in WT and VM hosts, we stratified our analysis into the CD4 and CD8 T cell compartments. Within the WT-derived CD4 subset, a population of CD44+ CD62L-CD127+ CD69-effector memory CD4 T cells was enriched in the VM host but not the WT host; OTII T cells with this phenotype were not found in either host (**Figure 4E**). Similarly, within the WT-derived CD8 subset, a population of CD44+ CD62L-PD-1+ CD69-effector T cells was enriched in the VM host but not the WT host; OTI T cells with this phenotype were not found in either host (**Figure 4F**).

To expand upon our flow cytometric analysis on T cell activation using a complementary approach, we again utilized dimensional reduction (Figure S2A), unsupervised clustering, and manual cluster annotation (Figure S2B, Table 2). Our UMAP was organized such that UMAP1^lo^ associated with CD4+ T cells, and UMAP1^hi^ associated with CD8+ T cells. UMAP2^lo^ associated with activated T cell markers CD44 and CD69 and UMAP2^hi^ associated with the naïve T cell marker CD62L (Figure S2C). Interestingly, while mice receiving WT donor BM showed a roughly expected 2:1 ratio of CD4 to CD8 T cells, mice receiving OT Tg donor BM instead showed a roughly 1:1 ratio of CD4 to CD8 T cells (Figure S2D). Consistent with our conventional flow cytometric analysis above, we found that that only WT donor T cells in both WT and VM host lung were mainly UMAP2^lo^, indicating an enrichment for effector T cell subsets (Figure S2D); whereas, OT Tg T cells in the lungs of both WT and VM hosts were predominately UMAP2^hi^, indicative of naïve and central memory T cells (Figure S2D). Additionally, we identified clusters corresponding to effector memory CD4 T cells and PD-1+ effector CD8 T cells, as indicated by red and lime green circles in our UMAP (Figure S2B) and confirmed their enrichment in WT→VM chimeras compared to WT→WT chimeras, as well as their near absence in OT donor T cells in either host (Figure S2D). Thus, Tg T cells can be recruited to the lung where they retain their naïve status despite significant concomitant tissue inflammation.

**Table 2.**
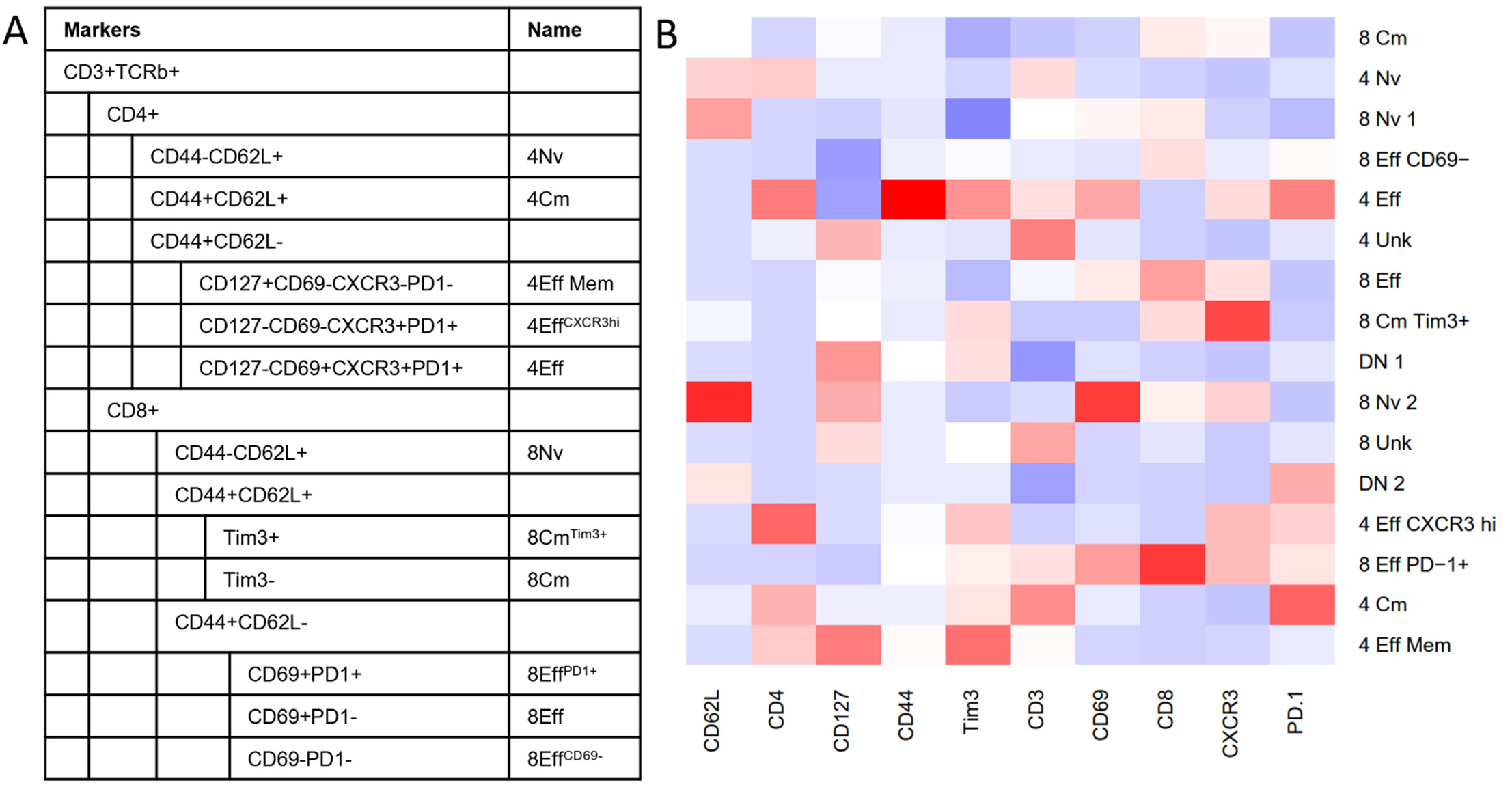
Annotation of T cell clusters by marker expression. (A) Manual annotation of 12 clusters identified by Phenograph unsupervised clustering of Live CD45+CD45IV-CD3+TCRβ+ lung extravascular (EV) αβ T cell flow cytometry events from WT+OTI/II→WT (n=5) and WT+OTI/II→VM (n=5) mice concatenated after excluding CD45.1 single positive radioresistant events.(B) shows a column normalized heatmap of median marker expression for clusters described in (A) and Figure S2B. Blue represents lower expression, white represents intermediate expression, red indicates higher expression.

**Figure S2.**
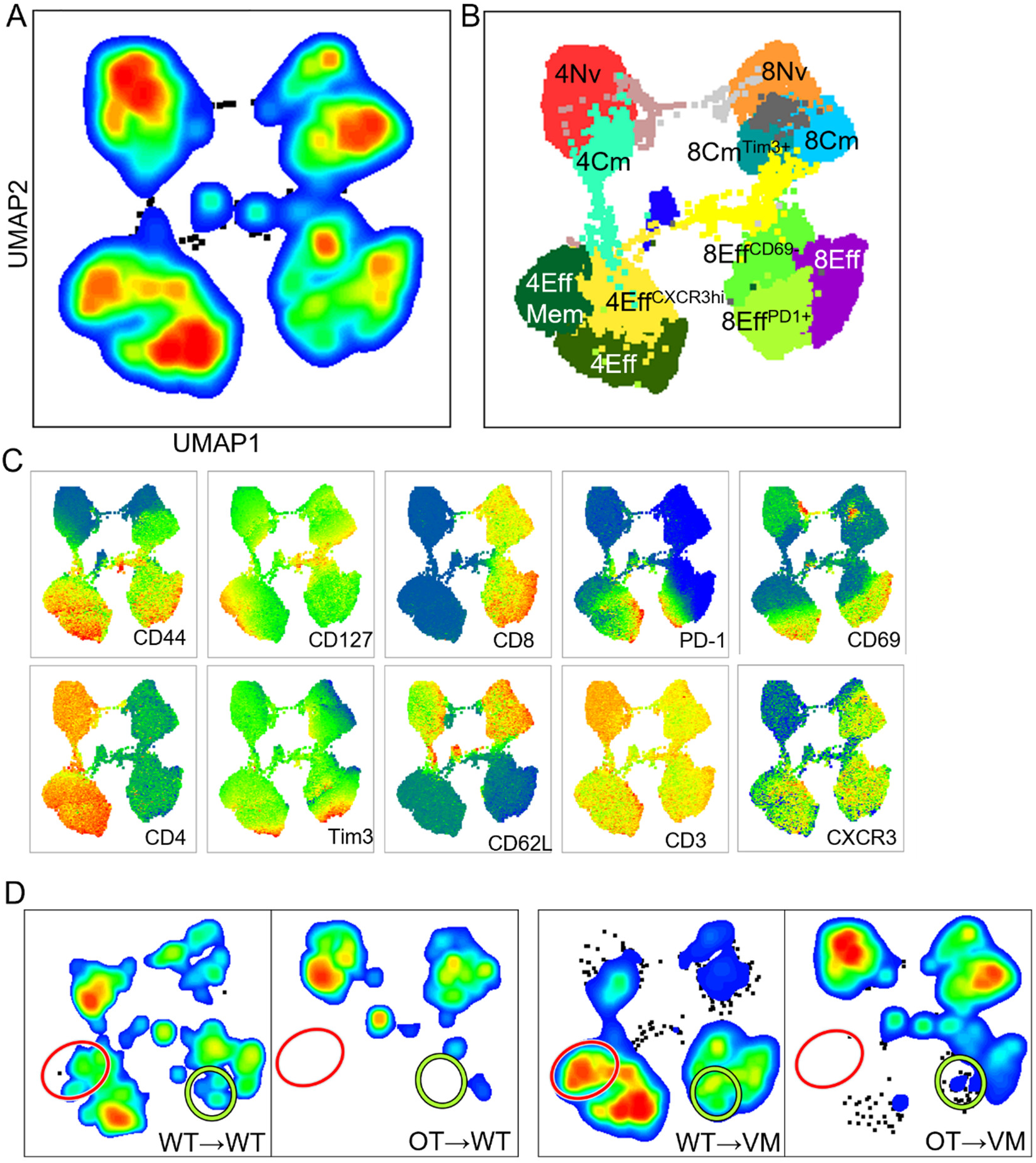
Unsupervised clustering of T cell markers identifies repertoire and VM host dependent phenotypes. 2000 Live CD45+ CD3+ TCRβ+ lung extravascular (EV) αβ T cell flow cytometry events per mouse from WT+OTI + OTII→WT (n=5) and WT+ OTI + OTII→VM (n=5) mice were concatenated after excluding CD45.1 single positive radioresistant events. (**A**) A UMAP was generated using CD44, CD127, CD8, PD-1, CD69, CD4, Tim3, CD62L, CD3, and CXCR3. (**B**) Unsupervised clustering using Phenograph identified 12 clusters in this dataset, which were manually annotated based on their expression profile (Table 2) and projected onto the UMAP alongside their annotated name. (**C**) Heatmap projections for each parameter used to generate the UMAP and perform unsupervised clustering. (**D**) UMAPs depicting events from WT donor-derived cells in either WT (WT→WT) or VM (WT→VM) hosts, as well as events from OTI/II donor-derived cells in either WT (OT→WT) or VM (OT→WT) hosts. Regions of the UMAP enriched in WT→VM events are highlighted with rings corresponding to the CD4 effector memory (CD4 Eff Mem; 4Eff Mem) cluster (red), and the CD8 effector PD-1 (CD8 Eff PD1; 8Eff PD1+) cluster (lime-green).

### WT→VM mice produce autoantibodies reactive against lung tissue and common autoantigens

The repertoire dependence of GC and PC formation in the VM lungs supported the notion that B cell differentiation in the VM lung depends on the recognition of self antigen. To determine whether WT B cells activated in a VM lung make autoantibodies, we first screened for anti-nuclear antibodies (ANAs) using an immunofluorescent (IF) HEp2 staining assay (**Figure 5A,B**). Remarkably, sera from WT→VM mice, but not the parental VM mice make a robust IgM ANA; however, there was only a minimal IgG response. To further explore the specificity of these ANA-reactive antibodies, sera from WT, VM and chimeric mice were screened on an autoantigen array. Consistent with the ANA staining patterns, only IgM antibodies from the WT➔VM chimeras, and no other groups, showed extensive autoantigen reactivity, particularly against autoantigens common to scleroderma/systemic sclerosis, and SLE/Sjogren’s (**Figure 5C,D**).

**Figure 5.**
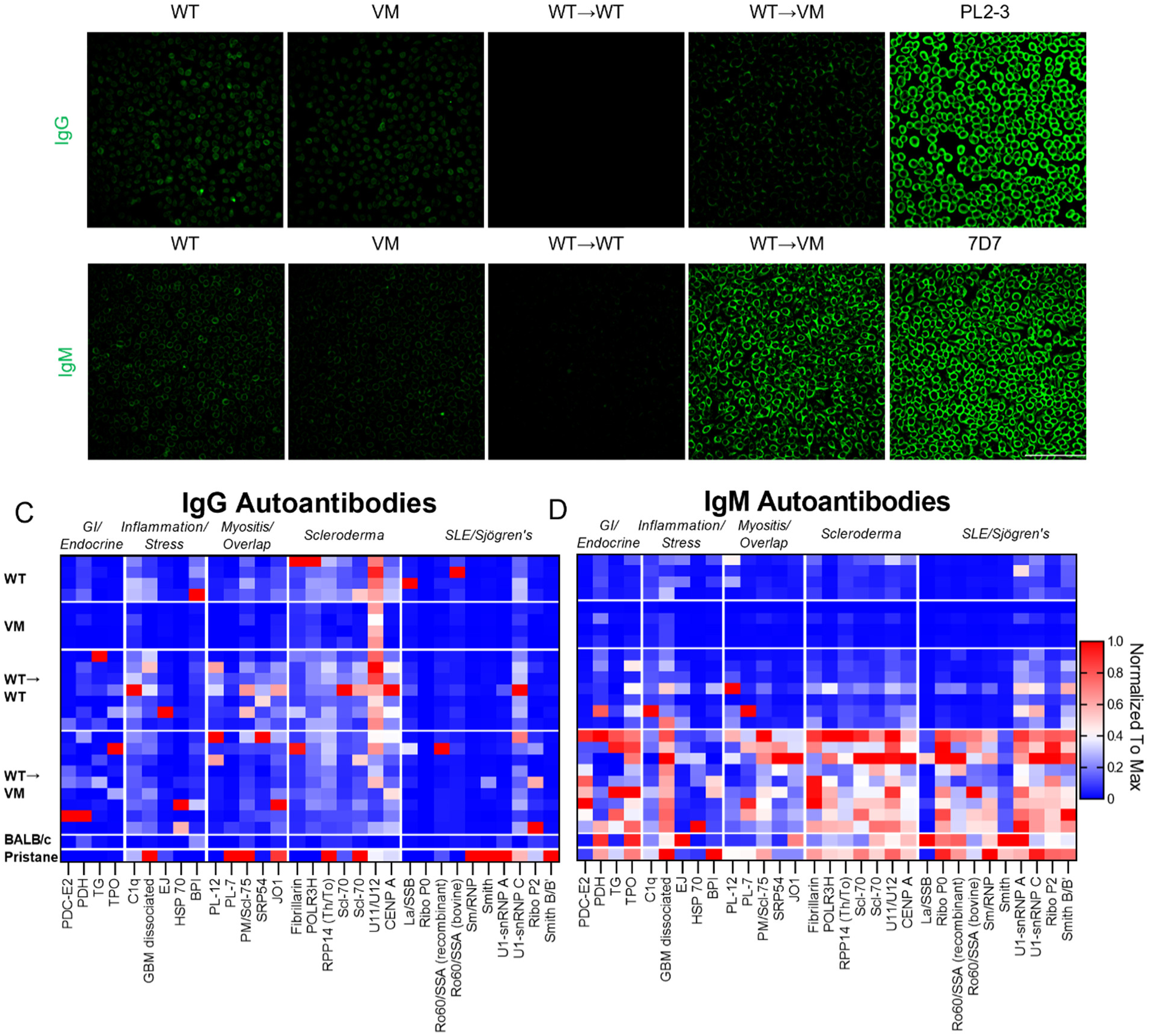
WT→VM mice produce IgM autoantibodies. (**A-B**) Immunofluorescent staining of HEp2 slides with representative sera of the indicated strains or chimeric mice with representative WT (n=5) and VM (n=3) mice, and WT→WT (n=12) and WT→VM (n=10) chimeric mice. (**A**) Autoantibodies were detected with a Dylight488 conjugated anti-IgG antibody (**A**) or anti-IgM antibody (**B)**. PL2-3 is an IgG2a anti-chromatin antibody and 7D7 is an IgM anti-DNA antibody (**C-D**) Sera from WT (n=4), VM (n=4), WT→WT (n=7), WT→VM (n=9), BALB/c (n=1), and BALB/c mice treated with pristane (n=1) were assessed by an autoantigen array. Data are column normalized to the highest and lowest signal observed for each autoantigen. Autoantigens are grouped according to the disease they are associated with: gastroenteric and endocrine autoimmunity (GI/Endocrine), non-specific inflammation and stress, myositis and overlap syndromes, scleroderma and systemic sclerosis, and SLE/Sjogren’s (**C**) shows the reactivity of sera IgG, (**D**) shows the reactivity of sera IgM.

Given the sera auto-antibody profile of WT➔VM chimeras aligned with several systemic autoimmune diseases that frequently target the lung, we also screened sera for reactivity with lung tissue. We assessed direct binding to lung tissue by staining lung sections from Rag1-/- mice with chimeric sera and then detecting bound antibodies with fluorophore conjugated anti-mouse IgM antibodies and found that sera Ig from WT➔VM chimeras clearly stained lung tissues (**Figure 6A**). To confirm these findings using a complementary approach, lung lysates from Rag1-/- mice were used for western blot analysis. As anticipated, IgM (but not IgG) from WT➔VM, but not WT➔WT sera, detected a number of distinct protein bands, several of which were bound by more than one serum sample (**Figure 6B,C**).

**Figure 6.**
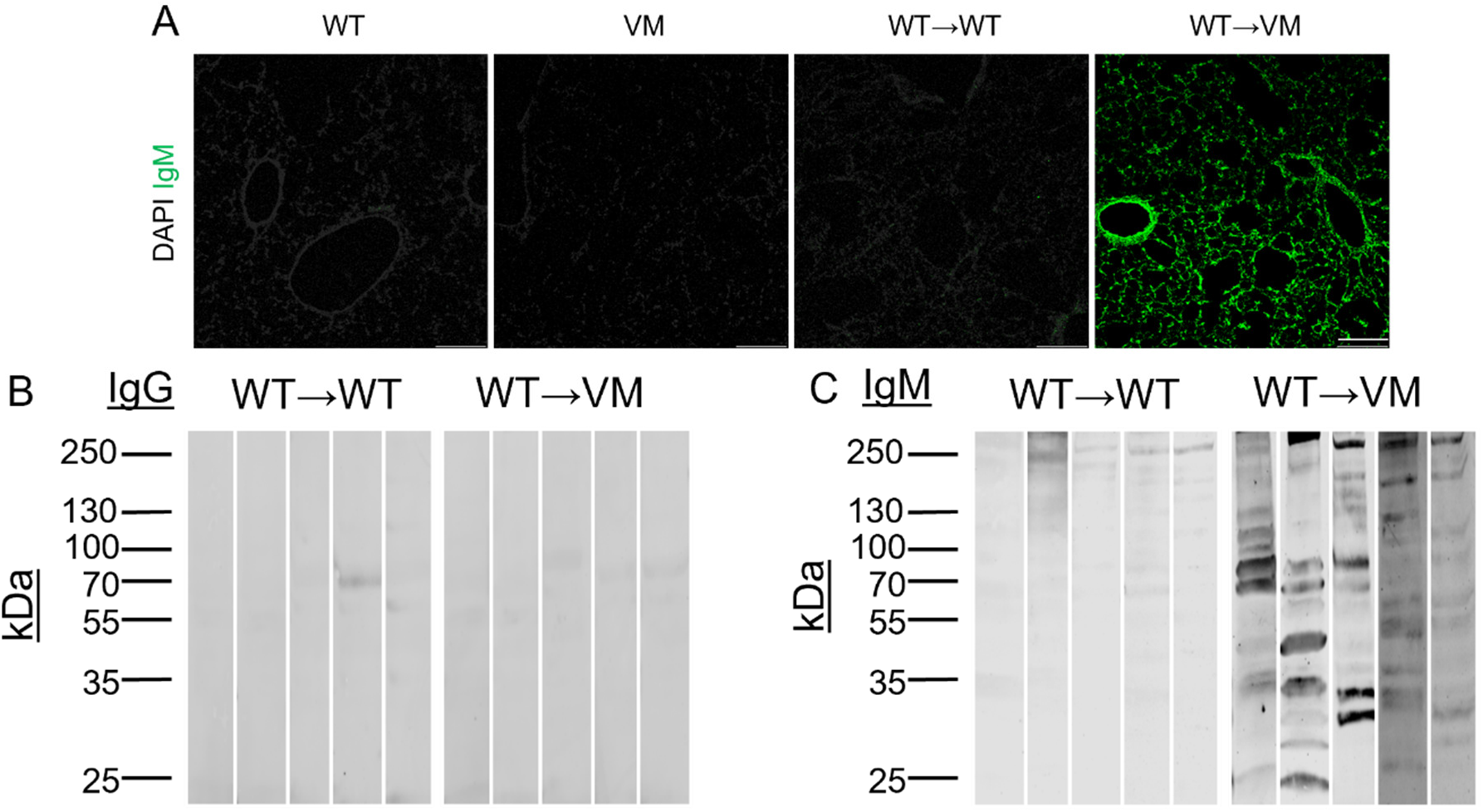
WT→VM autoantibodies target lung proteins. (**A**) Immunofluorescent staining of Rag1^-/-^ lung sections with serum samples diluted 1:40. A representative result is shown from WT (n=3) and VM (n=2) mice, and WT→WT (n=10) and WT→VM (n=10) chimeric mice. Slides were then developed with DAPI and AF488 conjugated anti-IgM antibody. Scale bars are representative of 200µm. (**B-C**) Lung reactive (**B**) IgG and (**C**) IgM antibodies were detected in sera from WT→WT or WT→VM chimeric mice (diluted 1:500) by western blot against lysates from Rag1^-/-^ lungs. Numbers on the left-hand side correspond to the mass in kDa of bands from the protein ladder. Data are representative of a total of (n=9) biologic replicates per group.

The development of autoantibodies in other models of STING-dependent autoinflammation has been shown to be partially dependent on T cells^23^. To determine if T cells might also contribute to the development of autoantibodies in our WT➔VM chimeric mice, we made radiation chimeras using BM from TCRβ-/- TCRδ-/- donors. As anticipated, at 8-10 weeks of age, these T cell deficient chimeric mice showed sera IgM and IgG that were both weakly autoreactive to HEp2 cells, Rag1-/- lung tissues, and lung lysates (Figure S5A-E). These data support a role for T cells in enhancing the development of lung autoreactive IgM antibodies in WT➔VM chimeric mice.

**Figure S3.**
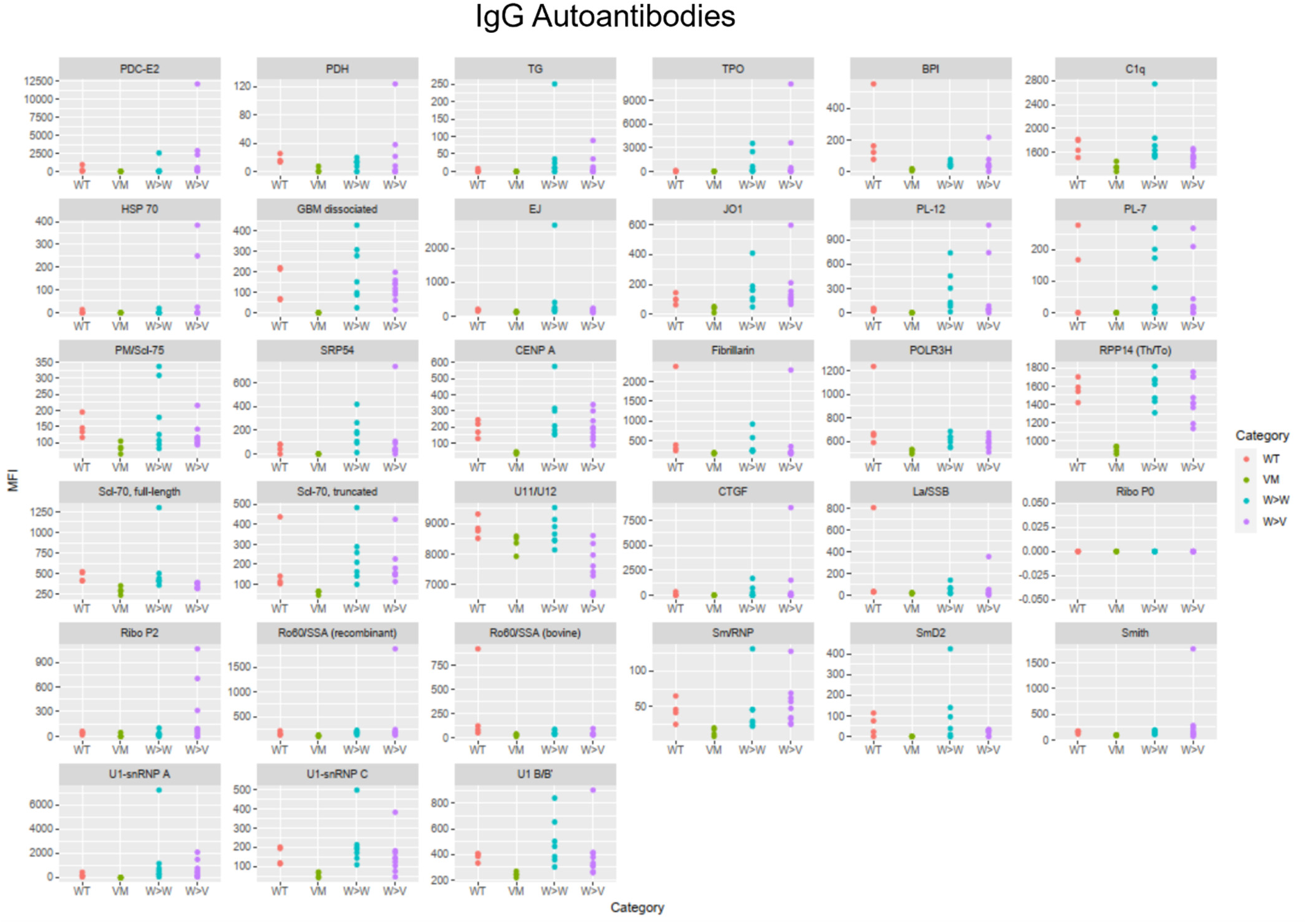
IgG Autoantigen Array. Autoantigen mean fluorescence intensity (MFI) of mouse sera IgG antibodies from WT (n=4), VM (n=4), WT→WT (n=7), WT→VM (n=9), BALB/c (n=1), and BALB/c mice treated with pristane (n=1) as detected using an anti-mouse IgG detecting antibody.

**Figure S4.**
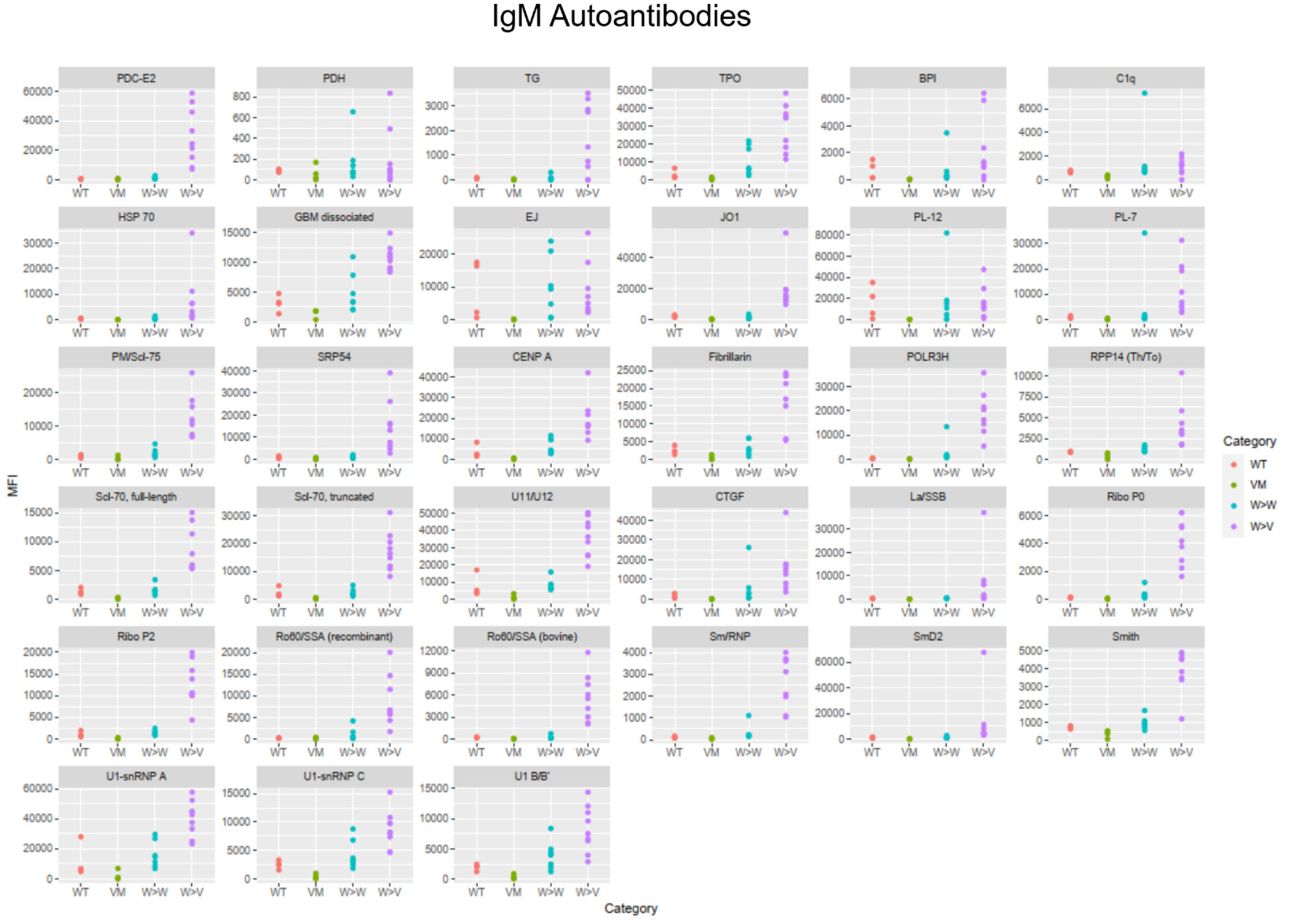
IgM Autoantigen Array. Autoantigen mean fluorescence intensity (MFI) of mouse sera IgG antibodies from WT (n=4), VM (n=4), WT→WT (n=7), WT→VM (n=9), BALB/c (n=1), and BALB/c mice treated with pristane (n=1) as detected using an anti-mouse IgG detecting antibody.

**Figure S5.**
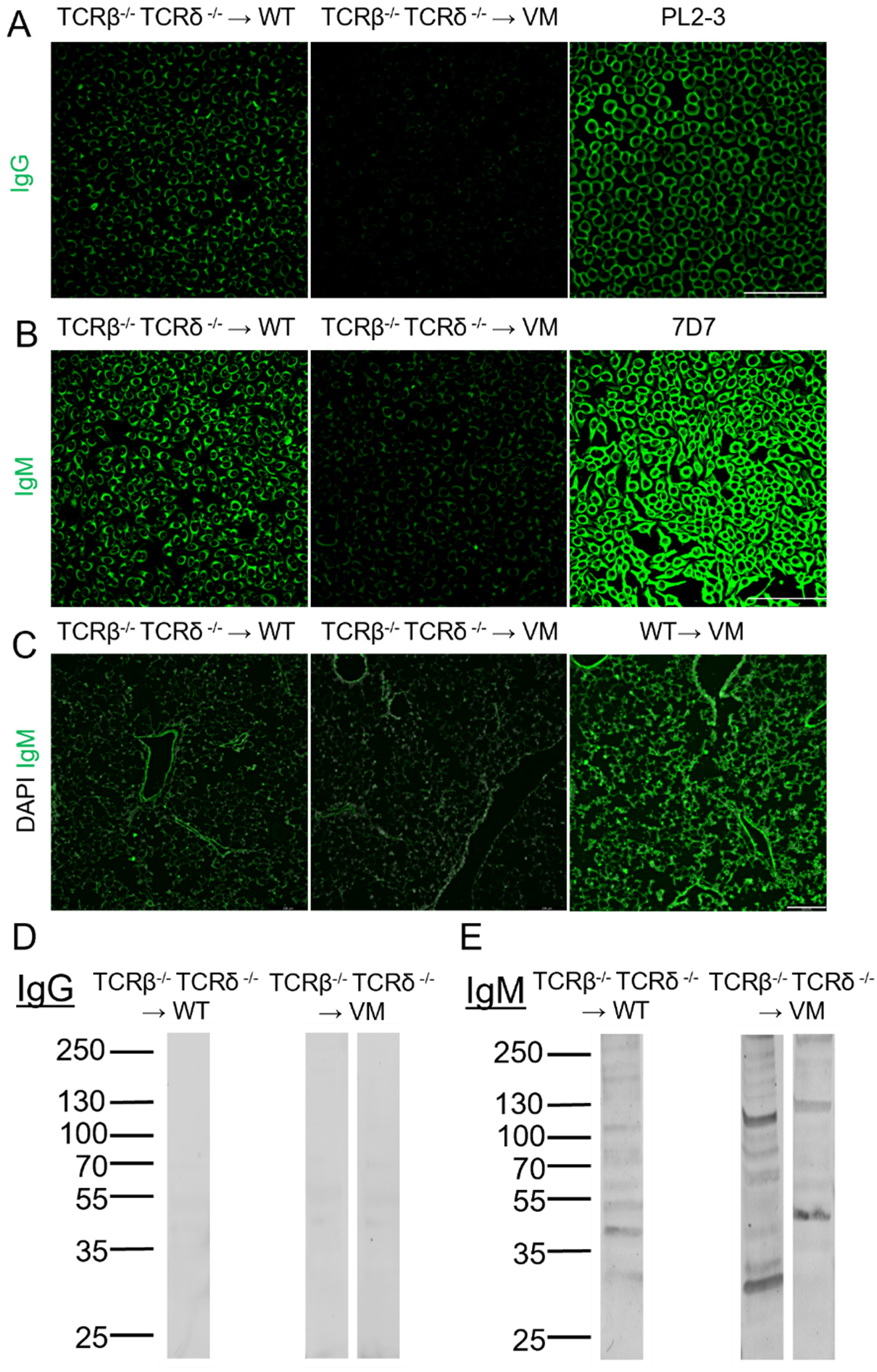
T cells contribute to autoantibody production in WT➔VM chimeras. (**A-B**) Immunofluorescent staining of HEp2 slides with 1:50 diluted sera. A representative result is shown from TCRβ-/- TCRδ-/- →WT (n=1) and TCRβ-/- TCRδ-/- →VM (n=2) chimeric mice. (**A**) Slides were then developed with a Dylight488 conjugated anti-IgG antibody (**B**) or anti-IgM antibody. PL2-3 is an anti-chromatin antibody, used as an IgG anti-nuclear antibody control, and 7D7 is an anti-DNA antibody, used as an IgM anti-nuclear antibody control. (**C**) Immunofluorescent staining of Rag1^-/-^ lung sections with 1:40 diluted sera. A representative result is shown from TCRβ-/- TCRδ-/- →WT (n=1),TCRβ-/- TCRδ-/- →VM (n=2), and WT→VM (n=10) chimeric mice. Slides were then developed with DAPI and AF488 conjugated anti-IgM antibody. Scale bars are representative of 200µm.

Altogether, these findings indicate that while VM mice produce very little circulating antibody, and show minimal antibody reactivity against self-antigens, autoreactive WT B cells that develop in VM hosts can differentiate into GC B cells and produce autoantibodies directed against specific self-antigens present on lung tissue.

### WT lymphocytes that develop in a VM host transfer ILD to a naïve Rag1KO mouse

Another hallmark of autoimmunity is the ability to adoptively transfer autoreactive lymphocytes into naïve mice and thereby transfer clinical manifestations of autoimmune disease^24^. Since lymphocytes in WT→VM chimeras showed a robust and functional autoimmune phenotype, we decided to test chimeric splenocytes for their capacity to transfer lung inflammation to naïve Rag1^-/-^ mice (**Figure 7A**). By 8 weeks post transfer, STING-sufficient Rag1-/- mice previously injected with splenocytes obtained from WT→VM, but not WT→WT, chimeric mice developed modest, histologically apparent immune aggregates in the lung (**Figure 7B**). IF staining revealed the presence of a CD3+ peri-broncho-vascular T cell infiltrate (**Figure 7C**). Furthermore, flow cytometric analysis of lung tissue from mice that received WT→VM splenocytes showed increased numbers of EV T cells compared to lungs from WT→WT splenocyte recipients (**Figure 7D**).

**Figure 7.**
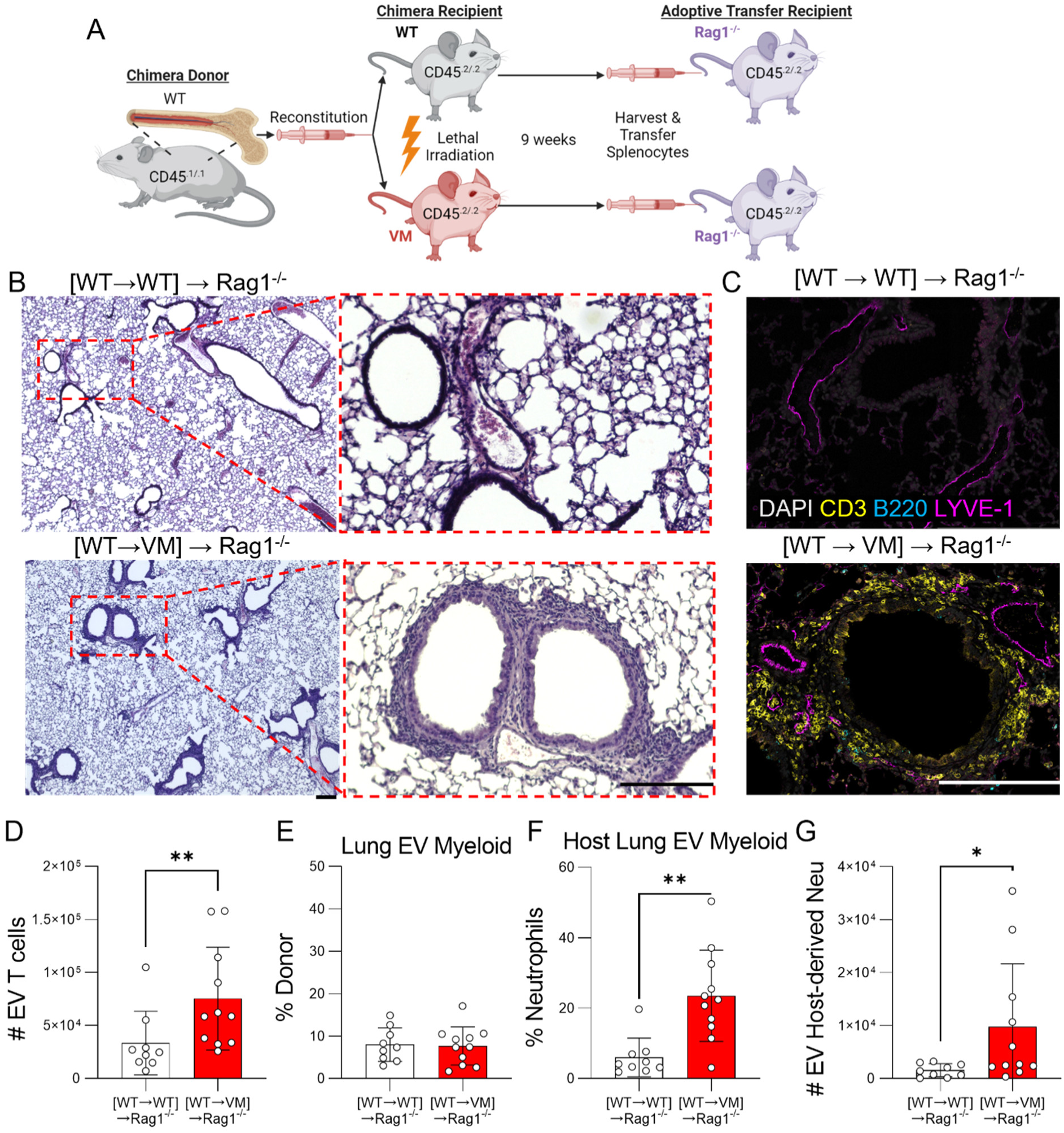
WT lymphocytes from WT➔VM chimeras induce lung inflammation in Rag1 KO mice. (**A**) 6-week-old CD45.2/.2 WT and VM mice were lethally irradiated and reconstituted with CD45.1/1 WT donor BM to generate WT→WT and WT→VM chimeric mice. Splenocytes from these chimeric mice were then harvested 8-weeks after BM engraftment and adoptively transferred into 8-week-old Rag1^-/-^ recipients to generate [WT→WT]→Rag1^-/-^ (n=8) and [WT→VM]→Rag1^-/-^ (n=11) mice and then evaluated 8-9 weeks later. (**B**) Representative H&E lung histology from [WT→WT]→Rag1^-/-^ and [WT→VM]→Rag1^-/-^ mice captured with a 4x objective. A region of interest, as indicated by dotted red box was further imaged with a 10x objective and shown approximately 5x magnified over the original image. (**C**) Lungs from [WT→WT]→Rag1^-/-^ and [WT→VM]→Rag1^-/-^ mice, stained for DAPI (gray), CD3 (yellow), B220 (cyan), and LYVE-1 (magenta). (**D**) Total number of EV T cells in the lungs of [WT→WT]→Rag1^-/-^ and [WT→VM]→Rag1^-/-^ mice. (**E**) Percentage of donor-derived cells in the CD11b+ or CD11c+ lung EV myeloid compartment. (**F**) Percentage and (**G**) number of CD11b+ Ly6G+ neutrophils from lung EV myeloid compartment. Nonparametric Mann-Whitney U-tests were used for pair-wise comparisons to determine statistical significance, (*p<0.05,**p<0.01). Scale bars in (B-C) are representative of 200μm.

One caveat of this adoptive transfer approach is that transfer of whole spleen also leads to transfer of donor myeloid cells. Thus, the effects we observe are not solely attributable to the transfer of a specific population of lymphocytes. However, we did not find any evidence for enrichment of donor myeloid cells in lungs from the recipients of WT➔VM chimeric cells (**Figure 7E**). However, Rag1^-/-^ host derived lung myeloid cells were significantly enriched for neutrophils in WT➔VM recipients compared to the WT—WT recipients by composition and absolute number (**Figure 7F, G**), consistent with previous experiments which demonstrated T cell-dependent recruitment of neutrophils to the lungs of VM mice^14^. Overall, these data suggest that the WT→VM chimera spleens contains a population of lung homing autoreactive effector T cells that can promote lung inflammation.

## Discussion

Autoinflammatory diseases driven by monogenic mutations affecting nucleic acid sensing are frequently associated with autoantibody production^4, 25^, and the development of autoantibodies is characteristic of numerous murine models of such monogenic diseases including Trex1-deficiency^23, 26^ and DnaseII-deficiency^27^. SAVI patients also commonly make autoantibodies^5^, although mice expressing the SAVI STING mutation V154M (VM) do not. By using WT➔VM radiation chimeras, we previously established that ILD in VM mice is driven by expression of the VM mutation in non-hematopoietic cells^14^. Using the same chimeric model, we now show that the inflammatory environment resulting from expression of the VM SAVI mutation by host-derived cells is sufficient to promote the activation of both autoreactive B and T cells, as well as the production of ANAs and antibodies reactive with proteins expressed in the lung. Thus, while lymphocyte-intrinsic STING activation constrains murine B cell activation and differentiation, VM-induced inflammation triggered by parenchymal/stromal cells in the lung promotes a break in tolerance and the development of autoimmunity in WT➔VM chimeras. Since human B cells do not express STING and therefore are not constrained by STING activation^13^, WT➔VM chimeras more closely recapitulate the development of autoantibodies as seen in human SAVI patients.

We found that autoantibodies produced by chimeric WT➔VM mice directly bind to lung tissues and show specificity for autoantigens commonly observed in scleroderma and Sjogren’s syndrome, autoimmune diseases which commonly present with interstitial lung disease^18, 28, 29^. WT➔VM mice develop increased numbers of germinal centers in both the spleen and BALT, suggesting that the airways may be a primary site of autoantigenic-driven responses. Intriguingly, most of the autoantibodies we detected in in the WT➔ VM chimeras were IgM, not IgG antibodies. Whether these antibodies are protective^30^ or pathogenic^14^ remains to be determined and will be explored in future studies.

In support of the lung as a primary site of tolerance break in SAVI disease, we have shown that although both WT and transgenic B cells expressing a BCR specific for a foreign (irrelevant) antigen both significantly accumulate in the lungs of VM mice, only non-transgenic B cells, with a diverse repertoire that includes self-reactive B cells, become sufficiently activated within the SAVI lung environment to contribute to a GC response. Similarly, the development of T cell effector subsets also requires a diverse repertoire. Moreover, T cells activated in a WT➔VM chimeric mouse retain the ability to home to lung tissues and promote neutrophil recruitment to the lung upon transfer to secondary unmanipulated naïve host. Thus, while SAVI disease initiates through excessive innate immune activation and autoinflammation, it evolves into an autoimmune process that incorporates autoreactive B and T cells.

The exact specificity of these lymphocytes remains to be determined. We expect that it will be possible to detect oligoclonal expansion of self-reactive B and T lymphocytes through BCR/TCR repertoire sequencing studies. In addition, future studies utilizing mass spectrometry to identify proteins repeatedly detected by western blot analysis of chimeric sera may help to identify specific proteins within lung tissues targeted by SAVI autoantibodies. This data may then be coupled to the identification of peptides that activate T cells present in the WT➔VM chimeras. Furthermore, although the lung is a major target of SAVI pathology, SAVI patients also present primarily with skin inflammation^5^, and SAVI mice have also been found to develop spontaneous colitis^31^. Thus, future studies may also seek to understand if these organ specific pathologies also feature an autoimmune component.

Overall, our data point to a break in peripheral tolerance in the SAVI lung. However, a previous report described a central tolerance defect in a murine model of COPA, another STING driven autoinflammatory disease which manifests with ILD. This study used radiation chimeras to show that excessive STING expression in non-hematopoietic thymic tissues led to a defect in negative selection and reduced numbers of regulatory T cells^32^. While central tolerance defects may also occur in WT➔VM chimeras, the fact that both MD4 B cells and OTI and OTII T cells are recruited to the VM lung to a greater extent than to a WT lung, indicates that additional factors beyond central tolerance defects drive SAVI ILD. Together these data provide evidence for a multi-step process wherein STING activation within non-hematopoietic lung cells promotes a repertoire-independent recruitment of immune cells into the lung environment, wherein repertoire-dependent activation of lymphocytes follows, resulting in the development of humoral autoimmunity directed against both lung tissues and systemic self-antigens.

Our findings build upon previous studies pointing to a role for T cell autoreactivity in SAVI disease. Wu et al. found that T cells expressing the SAVI STING mutation N153S (NS) were sensitive to ER stress driven apoptosis following TCR engagement. They further showed that crossing NS mice to OT-I mice rescued CD8 T cells from apoptosis and led to normal numbers of CD8 T cells in the spleen and peripheral blood of NS mice. Importantly, both systemic and lung inflammation were markedly reduced NS x OT-I mice^10^. The authors concluded that repertoire unrestricted CD8 T cells were required for SAVI ILD. Analysis of this data was limited by several factors. First, their study only assesses splenic lymphocytes, and did not directly examine the phenotype of the residual lymphocytes that still accumulated in the lungs of NS x OT-I mice. Secondly, while OT-I mice have normal numbers of CD8 cells, they are simultaneously deficient in CD4 T cells. Thus, the resulting phenotype of their NS x OT-I mice cannot be solely attributed to a limited CD8 T cell repertoire and may also reflect the absence CD4 T cell effector populations or other pathogenic factors produced by CD4 T cell subsets. The mixed chimera approach utilized in this manuscript resolves the limitations of Wu et al.’s previous study by building upon our observation that non-hematopoietic expression of the VM mutation is sufficient to initiate lymphocyte predominant lung inflammation^15^. This allowed us to directly compare the recruitment and activation of repertoire restricted CD4 and CD8 Tg T cells to repertoire-unrestricted WT cells, in the context of ongoing inflammation, and thus directly assess the contribution of both TCR and BCR repertoires.

It is also important to note that activated, and presumably self-reactive T cells and B cells, accumulate in the lungs of parental VM mice despite peripheral lymphopenia^15^. This indicates that the immune environment of the VM lung is sufficient to rescue receptor-activated VM lymphocytes from STING-induced apoptosis. The environmental cues that promote lymphocyte survival in these inflamed lungs warrant further investigation.

The results of our study have direct implications for the treatment of SAVI patients. We have confirmed that lymphocyte is initiated by STING activation in non-hematopoietic cells. We have further shown that VM expression in non-hematopoietic cells can trigger the activation of autoreactive T lymphocytes that can subsequently transfer lung pathology to a naïve host. It follows that inhibition of STING in patients with ongoing disease may not prevent continued pathology caused by existing lymphocytic effector mechanisms. Thus, therapeutic approaches which mitigate both the innate immune activity of non-hematopoietic VM parenchymal cells as well as existing autoimmune lymphocytes may be required for the effective treatment of SAVI patients.

## Materials and Methods

### Mice

STING V154M SAVI mice were generated as described^6^. SAVI VM mice and WT littermate controls were generated by crossing heterozygous male mice with WT females. TCRβ KO TCRδ KO (Jax #002122), μMT KO (Jax #002288), MD4 (Jax #002595), and B6 CD45.1 (Jax #002014) mice were obtained from The Jackson Laboratory. MD4 mice were backcrossed to Rag1 KO mice (Jax #002216) to generate MD4 Rag1 KO mice. OT-I Rag2 KO (Taconic #2334) and OT-II Rag2 KO (Taconic #11490) mice were obtained from Taconic Biosciences. VM mice were also intercrossed with B6 CD45.1 mice to generate CD45 allotype distinct hosts for chimera experiments. All mice used in these experiments were maintained in the same room and racks. STING^VM/WT^ (VM) and STING^WT/WT^(WT) littermate and sex-matched controls were used for all experiments. Roughly equal numbers of male and female mice were used. Serum was collected by cardiac puncture of euthanized animals. All animal experiments were conducted in accordance with the Institutional Animal Care and Use Committees at the University of Massachusetts Chan Medical School.

### Genotyping

Mice were genotyped using a combination of in-house genotyping PCRs performed on ear-clip DNA isolated from mice during weaning and custom probe-based genotyping performed by Transnetyx using real-time PCR to detect the STING V154M mutation.

### Generation of Radiation Chimeras

Lethally irradiated (850R) 6- to 8-week-old mice were reconstituted with 10^7^ bone marrow cells from sex and age matched mice. Mice were then placed on sulfatrim water and evaluated 8 to 9 weeks later.

### Splenocyte adoptive transfer

Splenocyte adoptive transfer was performed as previously described^27^. Single cell suspensions were generated from donor spleens and RBC lysed (Sigma-Aldrich R7757). 30×10^6^ splenocytes were delivered into sex-matched 8-week-old naïve Rag1 KO recipient mice. Engraftment of lymphocytes was confirmed 3 weeks later by analysis of blood. Mice were assessed 8 to 9 weeks after adoptive transfer.

### Intravascular labelling

Circulating immune cells were identified as previously described^33^. In brief, mice were injected i.v. with 3 μg of fluorescently-labeled CD45 mAb and euthanized 3 minutes later. Lung was collected without subsequent perfusion and digested as described below.

### Lung Digestion for hematopoietic cells

To assess immune cells and endothelia from the lung left lobe of lung was digested using a GentleMACS lung digestion kit (Miltenyi Biotec #130-095-927). In brief, lung was intratracheally inflated with 1ml of GentleMACS lung digestion kit digestion buffer. Lungs were then incubated at 37 C and dissociated using a GentleMACS Octo Dissociator with Heaters (Miltenyi Biotec #130-095-937). Cells were then filtered through a 70-micron mesh filter, spun down at 300xg for 10 minutes, and treated with RBC lysis buffer (Sigma #R7757) before subsequent assessments.

### B cell magnetic bead purification

Spleen was first dissociated in HBSS with 2% fetal bovine sera between 2 frosted glass slides, pelleted by centrifugation, and RBC lysed. B220+ B cells were purified by magnetic particle selection (BD Biosciences 551513).

### B cell proliferation assays

B cell proliferation was measured as previously described^34^. Briefly, B cells were stained with VPD450 at 3.5uM (BD Biosciences 562158). Cells were stimulated with BLYS at 50ng/ml and with an anti-IgM F(ab’)2 at 15ug/ml. On day 3, cells were stained with TO-PRO prior to flow cytometric assessment.

### Flow Cytometry

Cells were incubated in CD16/32 (Biocell #BE0307) and stained with antibodies as documented in Tables 3-6. Samples were fixed using Fluorofix buffer (Biolegend #422101). Intracellular staining was performed using a BD Biosciences Fixation/Permeabilization kit (BD Biosciences #554714) after stimulating cells for 4 hours with Brefeldin A (Biolegend #420601). Absolute cell counts were determined using counting beads (Biolegend #424902). Cells were acquired on an Aurora (Cytek) or a ZE-5 (Biorad) cytometer and analyzed with FlowJo software.

**Table 3.**
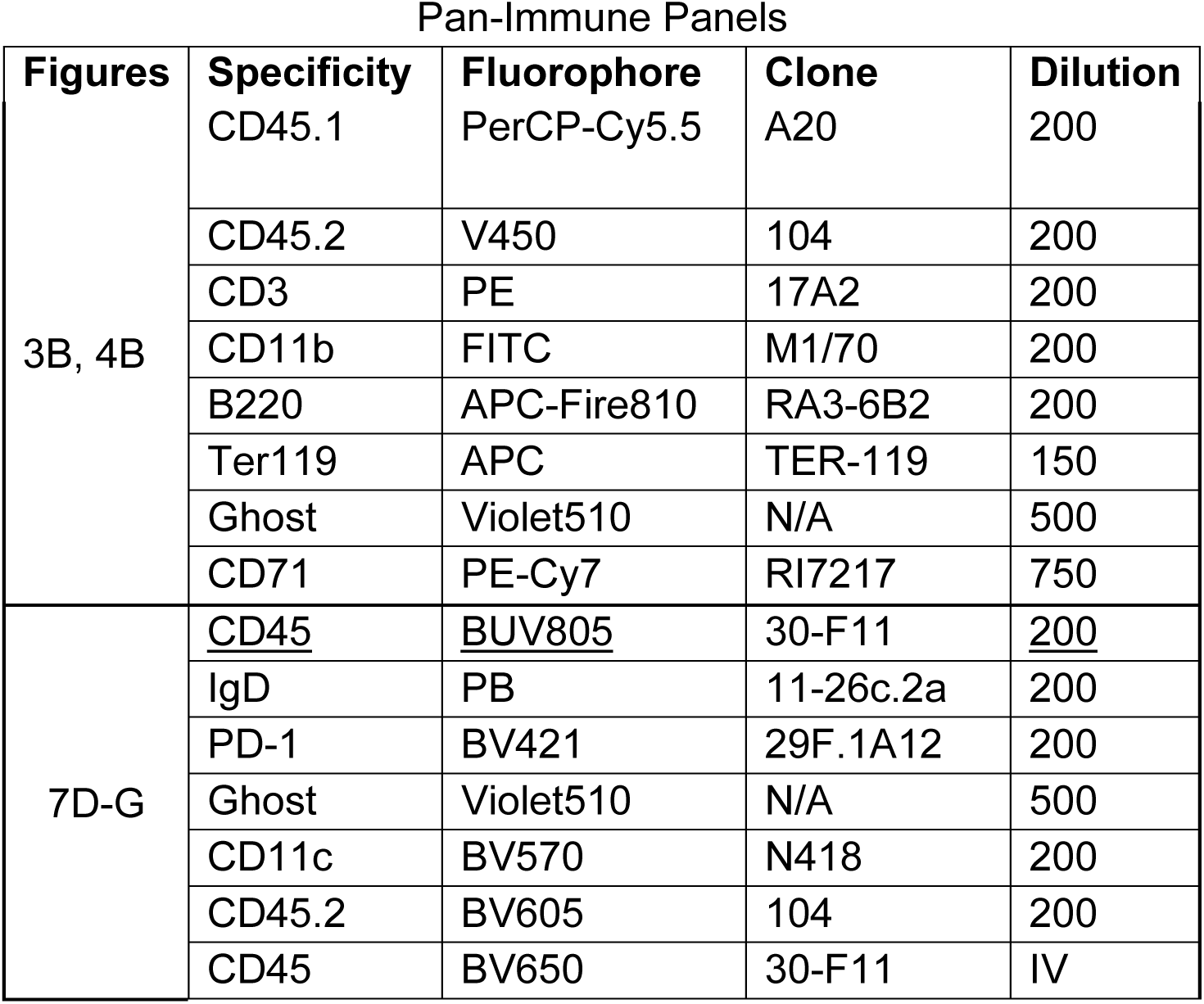
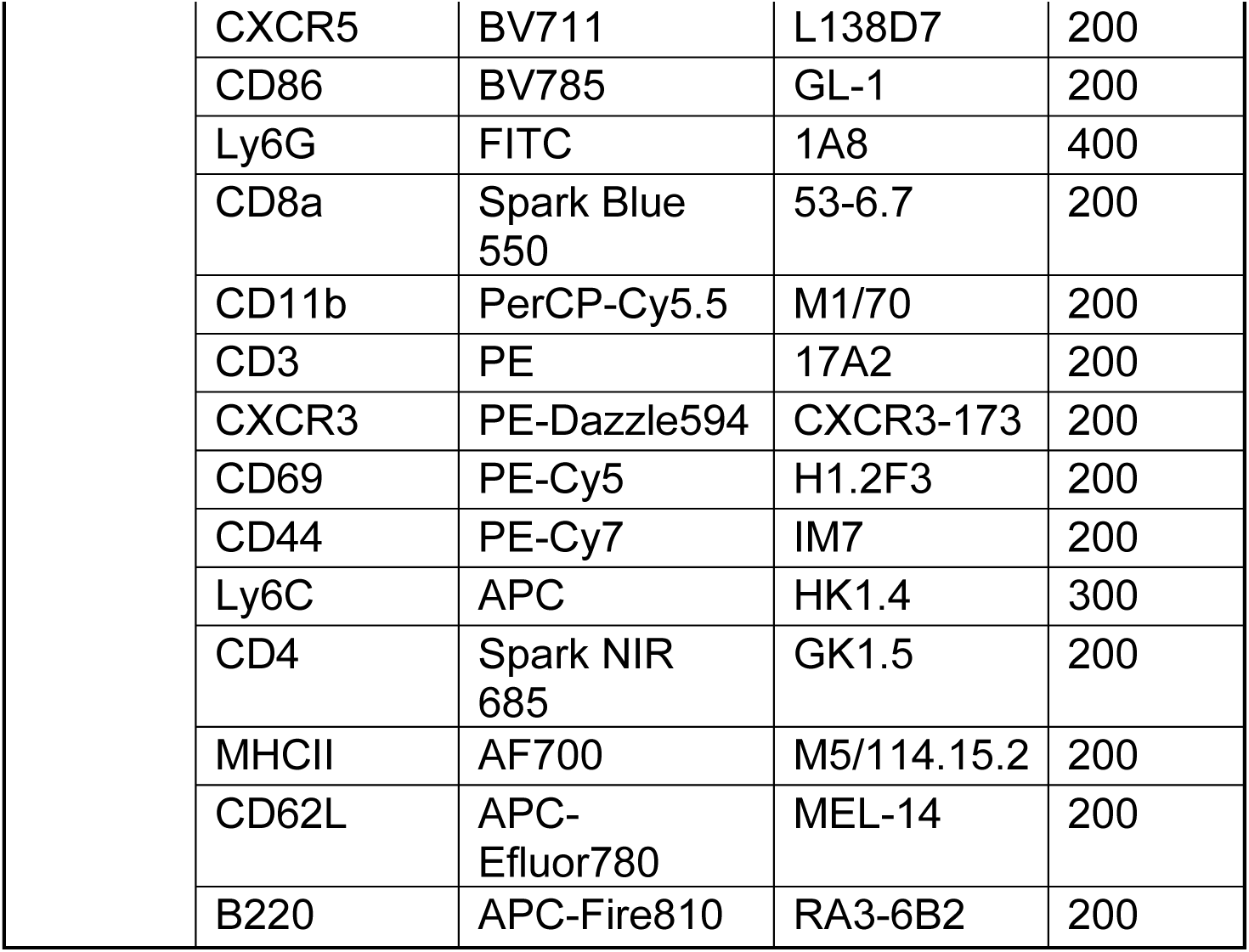
Pan-immune flow cytometry antibody panels.

| Pan-Immune Panels |  |  |  |  |
| --- | --- | --- | --- | --- |
| Figures | Specificity | Fluorophore | Clone | Dilution |
| 3B, 4B | CD45.1 | PerCP-Cy5.5 | A20 | 200 |
|  | CD45.2 | V450 | 104 | 200 |
|  | CD3 | PE | 17A2 | 200 |
|  | CD11b | FITC | M1/70 | 200 |
|  | B220 | APC-Fire810 | RA3-6B2 | 200 |
|  | Ter119 | APC | TER-119 | 150 |
|  | Ghost | Violet510 | N/A | 500 |
|  | CD71 | PE-Cy7 | RI7217 | 750 |
| 7D-G | <u>CD45</u> | <u>BUV805</u> | 30-F11 | <u>200</u> |
|  | IgD | PB | 11-26c.2a | 200 |
|  | PD-1 | BV421 | 29F.1A12 | 200 |
|  | Ghost | Violet510 | N/A | 500 |
|  | CD11c | BV570 | N418 | 200 |
|  | CD45.2 | BV605 | 104 | 200 |
|  | CD45 | BV650 | 30-F11 | IV |

|  |  |  |  |
| --- | --- | --- | --- |
| CXCR5 | BV711 | L138D7 | 200 |
| CD86 | BV785 | GL-1 | 200 |
| Ly6G | FITC | 1A8 | 400 |
| CD8a | Spark Blue 550 | 53-6.7 | 200 |
| CD11b | PerCP-Cy5.5 | M1/70 | 200 |
| CD3 | PE | 17A2 | 200 |
| CXCR3 | PE-Dazzle594 | CXCR3-173 | 200 |
| CD69 | PE-Cy5 | H1.2F3 | 200 |
| CD44 | PE-Cy7 | IM7 | 200 |
| Ly6C | APC | HK1.4 | 300 |
| CD4 | Spark NIR 685 | GK1.5 | 200 |
| MHCII | AF700 | M5/114.15.2 | 200 |
| CD62L | APC-Efluor780 | MEL-14 | 200 |
| B220 | APC-Fire810 | RA3-6B2 | 200 |

**Table 4.**
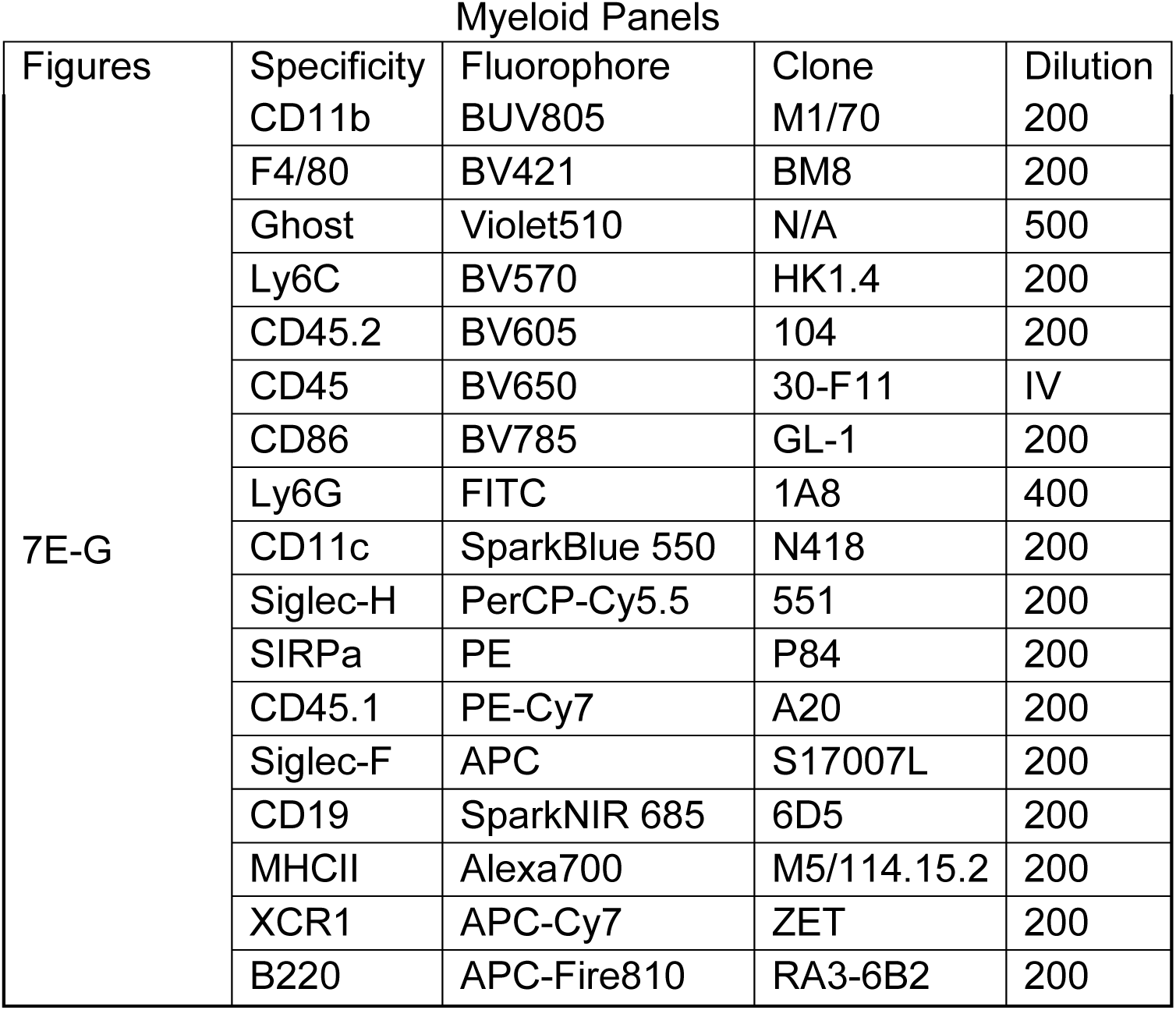
Myeloid flow cytometry antibody panels.

**Table 5.**
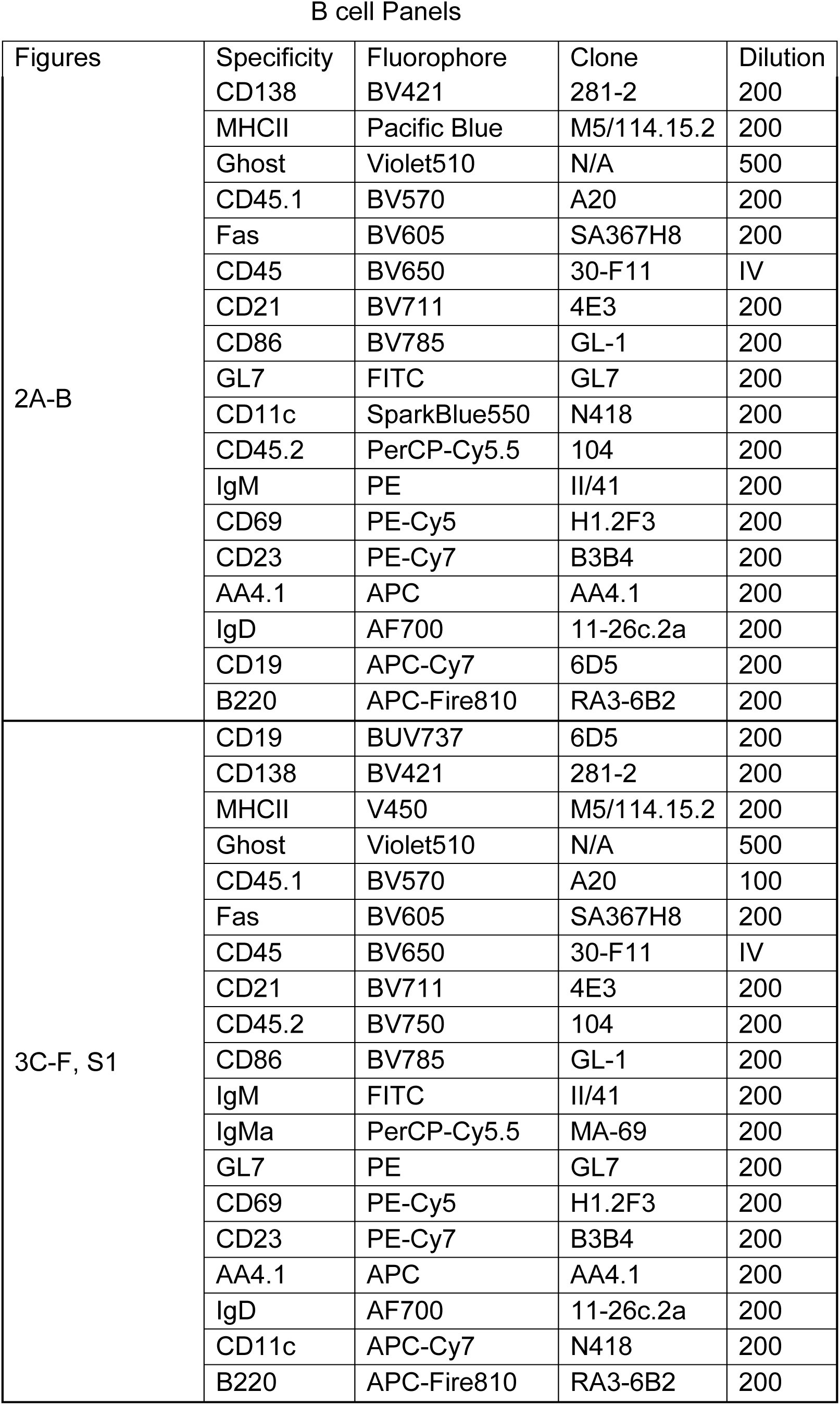
B cell flow cytometry antibody panels.

**Table 6.**
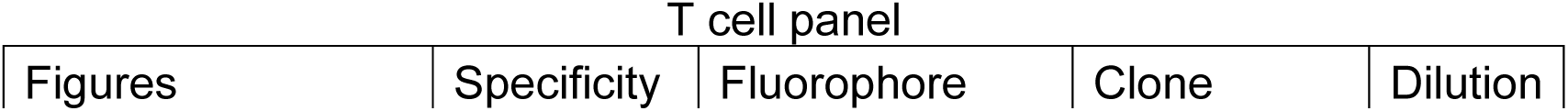
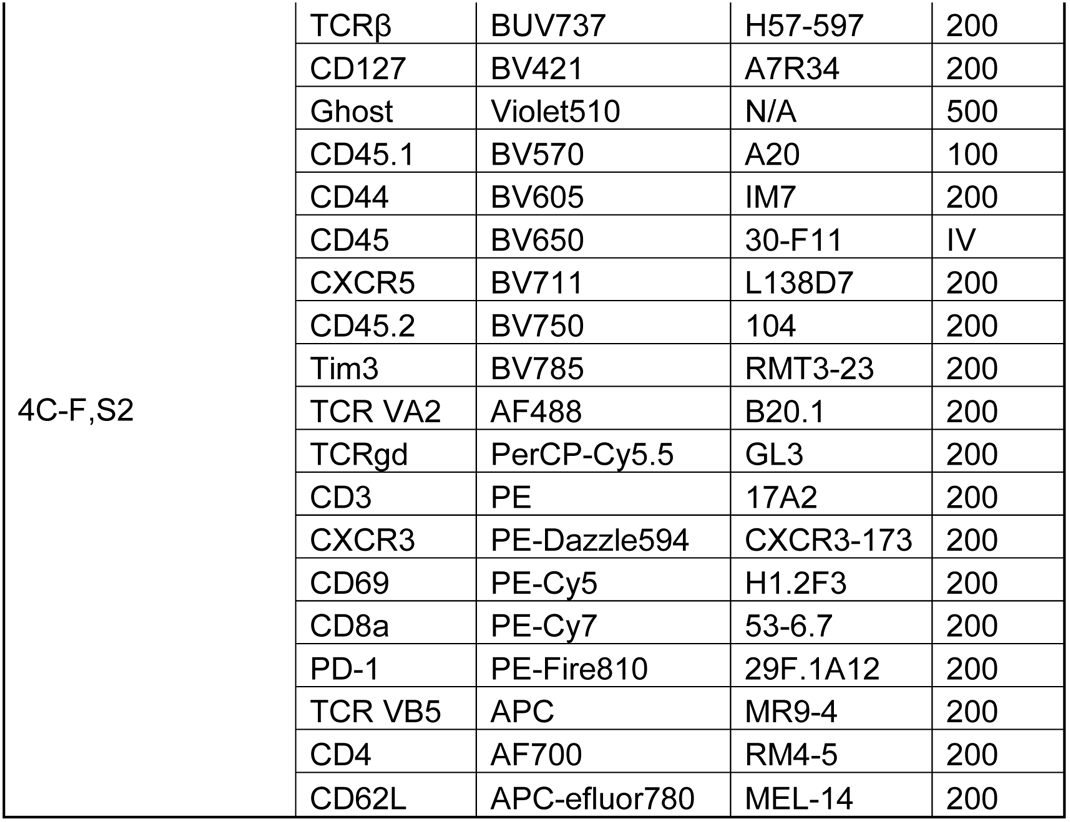
T cell flow cytometry antibody panels.

| T cell panel |  |  |  |  |
| --- | --- | --- | --- | --- |
| Figures | Specificity | Fluorophore | Clone | Dilution |
| 4C-F,S2 | TCRβ | BUV737 | H57-597 | 200 |
|  | CD127 | BV421 | A7R34 | 200 |
|  | Ghost | Violet510 | N/A | 500 |
|  | CD45.1 | BV570 | A20 | 100 |
|  | CD44 | BV605 | IM7 | 200 |
|  | CD45 | BV650 | 30-F11 | IV |
|  | CXCR5 | BV711 | L138D7 | 200 |
|  | CD45.2 | BV750 | 104 | 200 |
|  | Tim3 | BV785 | RMT3-23 | 200 |
|  | TCR VA2 | AF488 | B20.1 | 200 |
|  | TCRgd | PerCP-Cy5.5 | GL3 | 200 |
|  | CD3 | PE | 17A2 | 200 |
|  | CXCR3 | PE-Dazzle594 | CXCR3-173 | 200 |
|  | CD69 | PE-Cy5 | H1.2F3 | 200 |
|  | CD8a | PE-Cy7 | 53-6.7 | 200 |
|  | PD-1 | PE-Fire810 | 29F.1A12 | 200 |
|  | TCR VB5 | APC | MR9-4 | 200 |
|  | CD4 | AF700 | RM4-5 | 200 |
|  | CD62L | APC-eFluor780 | MEL-14 | 200 |

### Unsupervised Machine Learning Analysis of Flow Cytometric data

To analyze high-parameter flow cytometry data, we employed analysis by unsupervised machine learning tools. In brief, samples were annotated with meta-data to identify individuals within cohorts. Gating was performed to identify lineage positive populations across specific organ compartments (extravascular lung), and these populations were concatenated into a single FCS file. UMAP dimensional reduction was performed using select parameters for visualization. Unsupervised clustering was performed on the same parameters used for UMAP dimensional reduction using the Phenograph, as prior studies indicated that Phenograph showed high robustness with larger datasets^35^. Metadata annotations were then used to deconvolve concatenated data and re-segregate events into experimental and control groups. Cluster distributions for individuals within cohorts were calculated, and multiple comparisons testing was performed to identify differentially represented clusters across experimental conditions. The identity of clusters was determined through manual annotation using ClusterExplorer, conventional gating, and multi-parameter heatmap analysis. Analysis was performed in FlowJo 10 and Prism Graphpad 9.

### Immunoglobulin Analysis

ELISAs for IgM, IgG1, and IgG2a were measured using a previously described in-house sandwich ELISA^36^. Polyclonal goat anti-IgM (Southern Biotech 1020-01), anti-IgG1 (BD 557273), and anti-IgG2a (Southern Biotech 1077-01) capture antibodies were coated onto plates at 1ug/ml. A biotinylated-conjugated anti-kappa light chain antibody (BD 559750) and streptavidin-HRP (southern Biotech 1050-05) was used for detection.

### Histology

Lungs were dissected, inflated intratracheally with 10% phosphate buffered formalin (PBF) via a flexible catheter, fixed in 10% PBF at room temperature for 48 hours, and transferred into 70% ETOH. Lungs were then paraffin embedded, sectioned, and then stained with H&E by Applied Pathology Systems (Shrewsbury, MA). Whole H&E lung slides were scanned at 4x or 10x using an EVOS FL Auto microscope or an EVOS M7000 microscope housed in the Bone Analysis Core (University of Massachusetts Chan Medical School, Worcester, MA).

### Immunofluorescence

Sections were generated from either formalin fixed paraffin embedded (FFPE) or fresh-frozen lungs (OCT). For FFPE sections, 7 micron thick sections were prepared from FFPE blocks by the University of Massachusetts Chan Medical School Morphology Core. After deparaffinization using xylene, antigen retrieval was performed with 10mM Na Citrate 0.05% Tween 20 in a pressure cooker for 15 minutes. For OCT lung sections, lungs were inflated with a 1ml of a 1:1 PBS: OCT, tied off at the trachea with surgical suture, and snap frozen in a bath of dry ice chilled 70% ETOH. 7 to 40 micron thick sections were fixed in acetone for 10 minutes at −20C. FFPE or OCT sections were then permeabilized in 0.3% TritonX-100, blocked in 10% donkey sera, incubated with primary antibody overnight at 4C, incubated with secondary antibody for 1 hour at room temperature. Antibodies are listed in Table 8-9. Microscopy was captured on a Leica SP8 confocal microscope or a Leica Thunder widefield fluorescent microscope and analyzed in the Leica Application Suite X.

**Table 8.** Primary immunofluorescence antibody panels.

| Figures | Company | Clone | Host | Isotype | Target | Conjugate | Dilution |
| --- | --- | --- | --- | --- | --- | --- | --- |
| 7C | ABCAM | SP7/<br>ab16669 | Rabbit | IgG | CD3 |  | 150 |
|  | RND | Polyclonal/<br>AF2125 | Goat | IgG | LYVE-1 |  | 100 |
|  | BIO<br>LEGEND | RA3-6B2/<br>103204 | Rat | IgG | B220 | Biotin | 100 |
| 2C | BD | RA3-6B2/<br>550286 | Rat | IgG | B220 |  | 50 |
|  | ABCAM | RM1007/<br>ab281586 | Rabbit | IgG | CD45 |  | 1000 |
|  | VECTOR<br>LABS | B-1075-5 | Peanut | NA | PNA | Biotin | 250 |

**Table 9.** Secondary immunofluorescence antibodies.

| Figures | Company | Clone | Host | Isotype | Target | Conjugate | Dilution |
| --- | --- | --- | --- | --- | --- | --- | --- |
| 2C | Invitro<br>gen | Polyclonal | Goat | IgG | Rat<br>IgG | AF488 | 500 |
|  | Invitro<br>gen | Polyclonal | Goat | IgG | Rabbit<br>IgG | AF555<br>plus | 500 |
|  | Invitro<br>gen | N/A | Strept<br>avidin | IgG | Biotin | AF647 | 500 |
| 7C | Jackson | Polyclonal | Donkey | IgG | Rabbit IgG | AF488 plus | 500 |
|  | Invitrogen | Polyclonal | Donkey | IgG | Goat IgG | AF555 | 500 |
|  | Invitrogen | Polyclonal | Streptavidin | IgG | Biotin | AF647 | 500 |
| 5A-B, 6A | Jackson | Polyclonal | Goat | IgG | Mouse IgG | Dylight 549 | 300 |
|  | Invitrogen | Polyclonal | Goat | IgG | Mouse IgM | Dylight 488 | 300 |

### Anti-nuclear antibody assay

Slides coated with HEp-2 cells (Biorad #26102) were stained with sera diluted 1:25 in 0.2% BSA PBS and stained for 1 hour at room temperature. PL2-3 IgG was used as an IgG ANA positive control, 7D7 was used as an IgM ANA positive control and were both diluted 1:100. Slides were then washed and stained with Goat anti-mouse IgG-Dylight 488 (Invitrogen #A-11001) or Goat anti-mouse IgM-Dylight 488 (Invitrogen #SA5-10150) diluted 1:100 for 1 hour at room temperature, and then washed. Pro-long gold anti-fade mountant was applied and samples were visualized by fluorescent microscopy.

### Autoantigen Array

Autoantigen arrays were generated and assays on the arrays were performed as described previously^37^. 8 ug of each Ag, control mouse and anti-mouse antibodies were diluted in PBS and transferred into 96-well plates. Analytes were coupled to 1 × 106 carboxylated magnetic beads per ID (MagPlex-C, Luminex Corp.). Beads were distributed into 96-well plates (Greiner BioOne), washed, and re-suspended in phosphate buffer (0.1 M NaH2PO4, pH 6.2) on a 96-well plate magnet. Bead surface was activated by addition of 100 μL of phosphate buffer containing 0.5 mg of 1-ethyl-3(3-dimethylamino-propyl) carbodiimide (Pierce) and 0.5 mg N-hydroxysuccinimide (Pierce). After 20 min incubation on a shaker (Grant Bio), beads were washed and re-suspended in activation buffer (0.05 M MES, pH 5.0). Diluted Ags and control antibodies were incubated with beads for 2 hr at RT. The beads were washed 3 × in 100 μL storage buffer (0.02% PBS-T, 0.1% BSA, and 0.05% sodium azide), re-suspended in 50 μL storage buffer, and stored in plates at 4°C overnight. Immobilization of the Ags was confirmed, and the assay conditions were optimized by analysis of confirmed using commercially available mouse monoclonal antibodies or antibodies specific for 6X-histidine epitope tags (ab9108). Prototype human plasma samples were used for validation of bead arrays.

Mouse sera were diluted 1:100 in an assay buffer of 0.05% PBS-T supplemented with 3% (w/v) BSA (Sigma) and transferred into a 96-well plate. Negative assay control included BALB/cJ WT mouse serum and positive assay control included mouse serum from a pristane model of Systemic lupus erythematosus (SLE) as previously published^38^. The bead array was distributed into a 384-well plate (Greiner BioOne) by transfer of 5 μL bead array per well. 45 μL of the 1:100 diluted sera were aliquoted and transferred into the 384-well plate. Samples were incubated for 90 min on a shaker (Grant Bio) at RT. The beads were washed with 3 × with 60 μL PBS-T on a plate washer (EL406, Biotek). Secondary antibodies, including 50 μL of R-PE conjugated Fcg fragment specific goat anti-mouse IgG (Jackson), and R-PE conjugated Fcg fragment specific goat anti-mouse IgM (Jackson), were diluted 1:500 and 1:400 respectively in 3% BSA in 0.05% PBS-T based on optimized assay conditions and transferred to transferred to the two halves of the 384-well plate, allowing for a parallel detection of different isotypes. After incubation with the secondary antibody for 45 min, the plates were washed with 3 × with 60 μL PBS-T and re-suspended in 50 μL PBS-T for readout in a FlexMap3D instrument (Luminex Corp).

Data analysis and visualization was performed using R Studio (2023.03.0) and GraphPad Prism 9.5.1 (GraphPad Software, Inc., San Diego, CA). Data was normalized by subtracting MFI values of unconjugated bare beads from antigen-coupled beads for each sample^39^. The normalized datasets were used to generate heatmaps in R Studio using the “ggplot2” package. while heatmaps were generated with Prism.

### Rag1 KO lung lysate western blots

Lungs were dissected from Rag1 KO mice following cardiac perfusion with 7ml of ice cold PBS, followed by three intra-tracheal washes with 1ml of ice cold PBS, and placed into M Tubes (Miltenyi #130-093-236) with 6ml of RIPA Lysis buffer (Thermo Scientific #89900) with 1x HALT protease inhibitor cocktail (Thermo Scientific #78429). Lungs were disassociated with a gentleMACS octo-dissociator using the Protein_01 program. Samples were then incubated on ice for 30 minutes and then sonicated for 2 minutes using 10 second sonication, 10 second rest cycles. Samples were then centrifuged for 4000xg for 20 minutes at 4C. Lysate protein concentration was quantified using a DC assay (Biorad #3000111). Nupage reducing buffer and loading buffer was added to lysates prior to boiling samples at 95C for 5 minutes. 30ug of lysate protein was loaded into wells of a Mini-Protean Precast gels (Biorad, 4568024) alongside PageRuler Plus stained protein ladder (ThermoScientific #26619). SDS-PAGE was then performed at 120V for 1 hour. Protein was then transferred onto a nitrocellulose membrane using semi-dry transfer run at 110mA for 80 minutes. Membranes were then blocked with 5% non-fat dry milk in 0.1% TBS-Tween20. Membrane was then stained with 1:500 sera diluted in blocking buffer overnight at 4C with shaking. After washing off sera, secondary antibody cocktail was added containing 1:20,000 goat-anti mouse IgG IRDye 680RD (LI-COR #926-68070) and 1:20,000 goat-anti mouse IgM IRDye 800CW (LICOR #926-32280). Membranes were then visualized using a LI-COR Odyssey Imager and analyzed in Image Studio Lite V5.2.

### Statistical Analysis

Since we were unable to confirm that our data was normally distributed, we used non-parametric tests for all statistical analyses with n<15. For single comparisons, Mann-Whitney tests were performed. For one-way ANOVA, the Kruskal-Wallis test was used. For multiple comparisons testing, multiple-Mann-Whitney test was used.

## Acknowledgements

Sharon Subramanian, Xhuliana Picari, Kristin Pike, and Stephanie Moses assisted in mouse breeding, genotyping and other experimental procedures. Thank you to Dr. Andrea Reboldi constructive feedback and discussion. These studies were supported by: NIHAID R01 AI128358, and Lupus Research Alliance Innovation Award. KMG was supported by: National Heart, Lung, and Blood Institute (NHLBI) F30 HL154674, National Institute of General Medical Sciences (NIGMS) T32 GM107000, National Institute of Allergy and Infectious Diseases (NIHAID) T32 AI132152, We declare the following conflicts of interest: Kate Fitzgerald is a Scientific Founder of Danger Bio, a Related Sciences company and is a member of the Scientific Advisory Board for Vesigen Therapeutics, NodThera, Janssen and Generation Bio.

